# Astrocytic modulation of information processing by layer 5 pyramidal neurons of the mouse visual cortex

**DOI:** 10.1101/2020.07.09.190777

**Authors:** Dimitri Ryczko, Maroua Hanini-Daoud, Steven Condamine, Benjamin J. B. Bréant, Maxime Fougère, Roberto Araya, Arlette Kolta

## Abstract

The most complex cerebral functions are performed by the cortex which most important output is carried out by its layer 5 pyramidal neurons. Their firing reflects integration of sensory and contextual information that they receive. There is evidence that astrocytes influence cortical neurons firing through the release of gliotransmitters such as ATP, glutamate or GABA. These effects were described at the network and at the synaptic levels, but it is still unclear how astrocytes influence neurons input-output transfer function at the cellular level. Here, we used optogenetic tools coupled with electrophysiological, imaging and anatomical approaches to test whether and how astrocytic activation affected processing and integration of distal inputs to layer 5 pyramidal neurons (L5PN). We show that optogenetic activation of astrocytes near L5PN cell body prolonged firing induced by distal inputs to L5PN and potentiated their ability to trigger spikes. The observed astrocytic effects on L5PN firing involved glutamatergic transmission to some extent but relied on release of S100β, an astrocytic Ca^2+^-binding protein that decreases extracellular Ca^2+^ once released. This astrocyte-evoked decrease of extracellular Ca^2+^ elicited firing mediated by activation of Nav1.6 channels. Our findings suggest that astrocytes contribute to the cortical fundamental computational operations by controlling the extracellular ionic environment.

**Key Points Summary:**

Integration of inputs along the dendritic tree of layer 5 pyramidal neurons is an essential operation as these cells represent the most important output carrier of the cerebral cortex. However, the contribution of astrocytes, a type of glial cell to these operations is poorly documented.
Here we found that optogenetic activation of astrocytes in the vicinity of layer 5 in the mouse primary visual cortex induce spiking in local pyramidal neurons through Nav1.6 ion channels and prolongs the responses elicited in these neurons by stimulation of their distal inputs in cortical layer 1.
This effect partially involved glutamatergic signalling but relied mostly on the astrocytic calcium-binding protein S100β, which regulates the concentration of calcium in the extracellular space around neurons.
These findings show that astrocytes contribute to the fundamental computational operations of the cortex by acting on the ionic environment of neurons.

## Introduction

The contribution of astrocytes to complex brain functions can no longer be overlooked. Multiple studies have linked astrocytic physiology to behavior or cognitive processes at the network and synaptic levels. However, their influence on information processing and integration at the cellular and circuit levels is still largely unresolved. Astrocytes are located in direct apposition and in between neurons (Araque et al. 2014), a strategic position to influence information processing and integration in the CNS. While this may not be the case in all parts of the CNS, at least in the cortex and some extent in the hippocampus, each astrocyte covers a well-defined territory with little overlap of neighboring astrocytic processes (Bushong et al. 2002). In rodent cortex, these processes are estimated to collect information from up to 90,000 synapses per astrocyte (Oberheim et al. 2006). Temporally, astrocytes work on a time scale of milliseconds to minutes, which enable them to integrate information over a much longer period of time than neurons (Araque et al. 2014). However, if their contribution to circuit operations is to be functionally relevant, astrocytes should respond selectively to stimuli that are proper to the circuitry in which they are embedded. This is indeed the case in sensory areas such as the barrel cortex (Wang et al. 2006), the olfactory bulb (Otsu et al. 2015; Petzold et al. 2008), the thalamic barreloid fields (Claus et al. 2018), the dorsal horn of the spinal cord (Sekiguchi et al. 2016), or the somatosensory (Ghosh et al. 2013; Schulz et al. 2012; Winship et al. 2007; Zhang et al. 2016) and visual cortices (Schummers et al. 2008). In the latter, presentation of visual stimuli elicits large Ca^2+^ responses in astrocytes which display sharper tuning curves than surrounding neurons (Perea et al. 2014; Schummers et al. 2008; Sonoda et al. 2018). This activity is likely to impact the circuit output since stimulation of astrocytes modulates orientation selectivity of interneurons and pyramidal neurons in layers 2 and 3 of the visual cortex (Perea et al. 2014). Here we examined at the cellular level whether astrocytic stimulation affected information integration and processing of layer 5 pyramidal neurons (L5PN).

L5PN convey the main output of the cortex (Bannister 2005; Thomson and Lamy 2007). These neurons have an elaborate dendritic arbor covering all cortical layers, with proximal – basal – dendrites receiving bottom-up (feedforward stream) inputs mostly from thalamus and local cortical neurons, and distal dendrites – the apical tuft – receiving top-down (feedback stream) connections from other cortical areas (Larkum 2013). Inputs directed to basal dendrites directly affect the generation of action potentials (AP), however, inputs directed to the apical tuft can affect AP output if there is a temporal association of backpropagating AP into apical dendrites (Stuart and Sakmann 1994) and a dendritic spike generated at the calcium (Ca^2+^) spike initiation zone located at the basis of the apical tuft (Larkum et al. 1999). Therefore, mechanisms governing integration of synaptic inputs in distal and proximal dendrites in L5PN may have significant consequences in cortical association, neuronal output, and ultimately behavior.

Here we used optogenetics to test whether astrocytic stimulation at the proximal level of the L5PN impacted processing of their distal inputs and found that it increased the ability of distal inputs to elicit firing and prolonged their effect on spiking activity. We characterized the cellular mechanisms involved.

## Materials and Methods

### Ethics statement

All procedures conformed to the guidelines of the Canadian Council on Animal Care and were approved by the animal care and use committees of the Université de Montréal (QC, Canada) and Université de Sherbrooke (QC, Canada). A total of 121 animals were used, including 25 wild-type mice (C57BL/6, Charles River), 20 mice expressing the genetically encoded Ca^2+^ indicator GCaMP6f (Chen et al. 2013b) under the control of the Thymus cell antigen 1 (Thy1) promoter, i.e. Thy1-GCaMP6f mice (C57BL/6J-Tg(Thy1-GCaMP6f)GP5.17Dkim/J, stock 25393, JAX), and 76 mice expressing the channelrhodopsin 2 (ChR2) under the control of the GFAP promoter (GFAP-ChR2) mice. GFAP-ChR2 mice were obtained by crossing GFAP-Cre (B6.Cg-Tg(Gfap-cre)73.12Mvs/J, stock 12886, JAX) with ChR2-lox mice (B6.Cg-Gt(ROSA)26Sortm32(CAG-COP4*H134R/EYFP)Hze/J, stock 24109, JAX). Sex of the individuals used was not considered in the present study. Care was taken to minimize the number of animals used and their suffering.

### V1 slice preparation

Coronal slices were prepared as previously described (Araya et al. 2006a,b, 2007, 2014) from 11 to 31-day old mice anesthetized with isoflurane and decapitated with a guillotine. The cranium was opened and the brain was removed and dipped in an ice-cold sucrose-based solution (in mM: 3 KCl, 1.25 KH_2_PO4, 4 MgSO_4_, 26 NaHCO_3_, 10 Dextrose, 0.2 CaCl_2_, 219 Sucrose, pH 7.3–7.4, 300-320 mOsmol/kg) bubbled with 95% O_2_ and 5% CO_2_. V1 coronal slices (350 μm thick) were prepared in the same solution with a VT1000S vibratome (Leica). Slices were then stored and allowed to rest at room temperature for an hour in a holding chamber filled with artificial cerebrospinal fluid (aCSF) (in mM: 124 NaCl, 3 KCl, 1.25 KH_2_PO_4_, 1.3 MgSO_4_, 26 NaHCO_3_, 10 Dextrose, and 1.2 CaCl_2_, pH 7.3–7.4, 290–300 mOsmol/kg) bubbled with 95% O_2_ and 5% CO_2_.

### Whole-cell patch-clamp recordings

Whole-cell recordings were carried out at room temperature in a recording chamber perfused with aCSF bubbled with 95% O_2_ and 5% CO_2_. Neurons and astrocytes were visualized under a FV1000 confocal microscope (Olympus) equipped with a 40× water-immersion objective and an infra-red CCD camera. Patch pipettes were pulled from borosilicate glass capillaries (1.5 mm outside diameter, 1.12 mm inside diameter; World Precision Instruments) using a P-97 puller (Sutter Instruments). For neuronal recordings, pipettes (resistance 6–8 MΩ) were filled with a solution containing (in mM) 140 K-gluconate, 5 NaCl, 2 MgCl_2_, 10 HEPES, 0.5 EGTA, 2 Tris ATP salt, 0.4 Tris GTP salt, pH 7.2–7.3, 280–300 mOsmol/kg, 0.05 Alexa Fluor 594 or 488, and 0.2% biocytin). Alexa Fluor 594 was used to monitor the presence of the large apical dendrite of the recorded pyramidal neuron during the experiment. After establishment of a gigaseal, the membrane potential was held at −60 mV, and the membrane patch was suctioned. The pipette resistance and capacitance were compensated electronically. Neurons were discarded when action potentials did not overshoot 0 mV or when the resting membrane potential was depolarized (>-45 mV). For astrocyte recordings, pipettes (resistance 4–6 MΩ) were filled with (in mM) 140 K-gluconate, 5 NaCl, 10 HEPES, 0.5 EGTA, 2 Tris ATP salt, 0.4 Tris GTP salt, pH 7.2–7.3, 280–300 mOsmol/kg and 0.2% biocytin. To identify astrocytes, Sulforhodamine 101 (SR101) labeling was performed in the holding chamber before the beginning of the experiment by adding to the aCSF solution 1 mM of SR101 for 20 min at 34°C, and then 20 min of wash out in aCSF (Condamine et al. 2018; Kafitz et al. 2008). Patched cells were considered astrocytes on the basis of their morphological characteristics (cell body ≤ ∼10 µm), and when showing a linear current increase following application of a 600 ms voltage ramp from −120 to +110 mV in voltage-clamp mode and no action potential following step current injections in current-clamp mode (Condamine et al. 2018; Panatier et al. 2011; Serrano et al. 2008). Additionally, astrocytes with a resting membrane potential above −60 mV in current clamp mode were rejected (Condamine et al. 2018). Whole-cell recordings were performed using a Multiclamp 700A amplifier and a Digidata 1322A digitizer coupled with a computer equipped with PClamp software (Axon Instruments).

### Immunohistochemistry

Coronal cortical sections (500 µm) were obtained from fresh brain tissue with a Vibratome VT 1000S (Leica), immersed immediately in 4% (wt/vol) paraformaldehyde in PBS and stored overnight at 4°C. Slices were then cryoprotected for 2h in 20% sucrose in PBS at 4°C and sectioned at 20 µm or 40-45 µm thickness with a sliding microtome (Leica). The tissues were then processed in function of the target immunofluorescence as described below.

To visualize cells expressing S100β, Nav 1.6 or GFAP, the sections were rinsed three times for 5 min with PBS and immersed for 60 min in a blocking solution containing 0.3% Triton X-100 and 10% normal goat serum in PBS. The sections were then rinsed in PBS three times for 5 min and incubated overnight at 4°C in the blocking solution, which contained the primary antibodies guinea-pig anti-S100β (Synaptic System 287004, lot 287004/3, dilution 1:400); and/or rabbit anti Nav1.6 (Alomone lab ASC-009, lot ASC009AN2302, dilution 1:400) and/or chicken anti-GFAP (Thermofisher Scientific AB4674, lot GR167007-2, dilution 1:400). The next day, the sections were rinsed three times for 5 min in PBS and incubated for 60 min in the blocking solution, which contained the secondary antibodies [goat anti-guinea-pig DyLight 594, Jackson Immunoresearch 106-515-003, lot 82831, dilution 1:500); goat anti-rabbit Alexa Fluor 488, Thermofisher Scientific A11034, lot 54533A, dilution 1:500); donkey anti-chicken Alexa Fluor 488, Jackson Immunoresearch 703-545-155, lot 12090, dilution 1:400)].

In our GFAP-ChR2-EYFP mice, the ChR2 is tagged with the enhanced yellow fluorescent protein (EYFP) which has largely the same structure as the green fluorescent protein (GFP) that can therefore be detected immunohistochemically with an anti-GFP antibody. To verify whether ChR2 was expressed in neurons, we combined immunodetection of EYFP with immunofluorescence experiments against the neuronal nuclei marker (NeuN) or microtubule-associated protein 2 (MAP2) which labels neuronal processes. To visualize cells expressing NeuN, EYFP or MAP2, the sections were first immersed for 30 min in a permeabilization solution that contained 0.4% Triton X-100. Then, they were immersed 60 min in a blocking solution containing 0.4% Triton X-100 and 5% bovine serum albumin in PBS. The sections were then incubated overnight at 4°C with the primary antibodies. The sections were then incubated overnight at 4°C with the primary antibodies (goat anti-GFP; Abcam AB5450, lot GR3319697, dilution 1:1000; rabbit anti-NeuN, Abcam AB104225, lot GR3321966, dilution 1:500; rabbit anti-MAP2, Abcam AB32454, lot GR3199625, dilution 1:500) diluted in 0.4% Triton X-100 and 1% bovine serum albumin in PBS. The next day, the sections were rinsed four times for 10 min in PBS and incubated for 90 min with the appropriate secondary antibodies (Donkey anti-goat Alexa Fluor 488, Abcam AB150129, lot GR3247118-1, dilution 1:500; donkey anti-rabbit, Alexa Fluor 594, Invitrogen A31572, lot 1837922, dilution 1:500). Biocytin was revealed by immersion in a PBS solution containing streptavidine-Alexa Fluor 594 (Molecular Probes S11227, dilution 1:400) for 60 to 240 min. In all cases, the sections were then rinsed again three to four times in PBS for 5-10 min and mounted on Fisherbrand microscope slides and cover slipped using Fluoromount-G (Southern Biotech) mounting medium.

Sections were observed and photographed using either an E600 epifluorescence microscope equipped with a DXM1200 digital camera (Nikon), an FV1000 confocal microscope (Olympus) or a TCS SP8 STED nanoscope (Leica). In all cases, removing the primary antibody resulted in the absence of specific labelling on brain sections. Photoshop CS5 (Adobe) was used to combine digital photomicrographs taken with different filter sets and to adjust the levels so that all fluorophores were clearly visible simultaneously.

### Optogenetics

In some experiments, the ChR2 was activated by using simultaneously a 440-nm laser and a 488-nm laser that were coupled with a FV1000 microscope (Olympus). The stimulation zone was delineated manually and limited to a small area around the recorded neuron and optogenetic stimulation was applied using 5-95 s light pulses (1-25% laser power). In some experiments, light was shone at other wavelengths using different lasers (559 or 589 nm) to determine whether the observed effects were specific to the light spectrum that activates the ChR2 (440-480 nm). In other experiments where very precise localisation of the stimulation zone was deemed to be less crucial, a Light-Emitting Diode (LED) system (Doric) was used to flash 20-60 s light pulses (intensity: 400-1000 mA) at 465 or 595 nm through an optic fiber (50 µm diameter) placed on the zone of interest at the surface of the brain slice.

### Ca^2+^ imaging

Rhod-2 AcetoxylMethyl (AM) (ThermoFisher Scientific) was used to record intracellular Ca^2+^ signals in GFAP-ChR2 mice because its excitation (552 nm)/ emission (581 nm) spectra do not interfere with ChR2 excitation spectrum (440-480 nm). For these experiments, the brain slice was placed at room temperature during 45 min in a chamber perfused with oxygenated aCSF containing 20 µM of Rhod-2-AM. The preparation was then installed under the microscope and perfused with aCSF as described for patch-clamp recordings. Changes in fluorescence were recorded using a FV1000 confocal microscope (Olympus) equipped with a set of appropriate filters (Morquette et al. 2015). To measure the changes in fluorescence, the regions of interest were delineated manually around the labelled cell bodies. Changes in fluorescence were acquired at a rate of ∼2 Hz and expressed as relative changes in fluorescence (ΔF/F) (Ryczko et al. 2016a,b, 2017). The baseline was defined as the averaged fluorescence before stimulation. Data analysis was carried out using ImageJ, Clampfit (Molecular Devices) and Matlab (Mathworks).

### Drugs

Chemicals were purchased from Sigma, Tocris Bioscience or Invitrogen. In some experiments, the Ca^2+^-binding protein S100β (129 µM) or the Ca^2+^-binding protein 1,2-bis(o-aminophenoxy)ethane-N,N,N′,N′-tetraacetic acid tetrasodium salt (BAPTA, 5 or 10 mM) were applied locally near the recorded L5PN. These applications were performed with glass micropipettes (tip diameter around 1 μm) using pressure pulses (2-20 psi) of variable duration (1-30 s) applied with a Picospritzer (Parker). In some experiments, the following drugs were bath applied: the specific Nav 1.6 channel blocker 4,9-anhydro-tétrodotoxine (4,9-TTX, 0.1 µM); the AMPA/kaïnate receptor antagonist 6-cyano-7-nitroquinoxaline-2,3-dione (CNQX, 10 μM), the NMDA receptor antagonist (2R)-amino-5-phosphonovaleric acid (AP5, 26 μM or 50 µM) and the GABA_A_ receptor antagonist gabazine (20 µM), the non-selective mGluR antagonist LY 341495 (100 µM) (Schoepp et al. 1999), and the purinergic receptor antagonist suramin (50 or 100 µM) (Torres et al. 2012). In some experiments, antibodies were applied as previously done (Condamine et al. 2018; Morquette et al. 2015) using larger pipettes (tip diameter 10-20 µm) that were carefully lowered to the surface of the tissue near the recorded cells. Each antibody of interest was slowly injected by applying a small pressure (0.1 psi) during 8-30 min before testing its effect on the recorded neuron. Two antibodies were used: a monoclonal mouse anti-S100β antibody (Sigma S2532, lot 123M4881, 40 μg/mL, sodium azide salt 140 μM) or a mouse anti-myosin heavy chain type IIA antibody as a control injection (Developmental Studies Hybridoma Bank SC-71, lot 4-23-15, 477 μg/ml, sodium azide salt 340 μM). S100β was obtained from Inixium (Laval, Québec, Canada) in aliquots at a concentration of 14.0 mg/mL supplemented with glycerol to a final concentration of 50% and stored at −80°C as previously described (Condamine et al. 2018). Before use, to eliminate the glycerol, aliquots of the protein were thawed and microdialysed overnight using a dialysis device (Float-A-Lyzer from Spectra/Por) floating in a 20 mM HEPES buffer containing 140 mM of NaCl at pH 7.4. The protein was then reconcentrated by centrifugation using centrifugal filter units (Amicon Ultra). The same buffer (20 mM HEPES containing 140 mM NaCl_2_, but no Ca^2+^) was used to suspend and adjust its concentration to 129 µM as previously described (Condamine et al. 2018). The number of moles of S100β were estimated by measuring the diameter of the droplet ejected at the tip of the pipette in the air following a single pressure pulse as previously described (Ryczko et al. 2016a,b, 2017). The droplet volume was estimated using the equation of a sphere and the number of moles was then calculated, as well as the mass taking into account a molecular weight of 10713 g/mol for S100β. The membrane potential of L5PN was maintained at the same value in control and drug conditions (see Table 2). This was done by monitoring the membrane potential and adjusting its value using current injection.

**Table 2.** Electrophysiological properties of the recorded neurons (mean ± sem) before and after drug application. The number of neurons is given between brackets. AB, antibody; Exp, experiment; LY, LY 341495; Opto, optogenetic activation of astrocytes; R, resistance; Sura, suramin; Vm, membrane potential. The number of neurons is given between brackets. **^a^** P > 0.05 vs. control, paired t test. **^b^** P > 0.05 vs. control, Wilcoxon Signed Rank test.

|  | Spike amplitude (mV) |  | Tau (ms) |  | R (MΩ) |  | Vm hold (mV) |  |
| --- | --- | --- | --- | --- | --- | --- | --- | --- |
| Exp. | Control | Drug | Control | Drug | Control | Drug | Control | Drug |
| Opto +<br>sura | 77.0 ±<br>3.9 | 77.8 ±<br>2.8<sup>b</sup><br>(n=4) | 38.8 ±<br>2.1 | 35.9 ±<br>8.0<sup>a</sup><br>(n=4) | 433.4 ±<br>100 | 278.8 ±<br>56.4<sup>b</sup><br>(n=4) | -57.3 ±<br>0.9 | -57.8 ±<br>1.1<sup>b</sup><br>(n=4) |
| Opto +<br>CNQX,<br>AP5,<br>gabazine | 84.6 ±<br>1.9 | 84.4 ±<br>1.9<sup>a</sup><br>(n=5) | 35.1 ±<br>6.8 | 45.5 ±<br>8.8<sup>b</sup><br>(n=5) | 259.4 ±<br>74.6 | 226.0 ±<br>60.9<sup>a</sup><br>(n=5) | -58.9 ±<br>1.0 | -59.0 ±<br>1.4<sup>a</sup><br>(n=7) |
| Opto +<br>CNQX,<br>AP5,<br>gabazine,<br>LY | 85.9 ±<br>2.2 | 84.4 ±<br>1.9<sup>a</sup><br>(n=4) | 43.1 ±<br>4.5 | 36.3 ±<br>6.7<sup>a</sup><br>(n=4) | 345.5 ±<br>43.1<sup>a</sup><br>(n=4) | 324.2 ±<br>73.9 | -56.8 ±<br>0.5 | -56.5 ±<br>0.6<sup>a</sup><br>(n=4) |
| Opto+<br>CNQX,<br>AP5,<br>gabazine,<br>LY, sura | 84.2 ±<br>4.2 | 85.2 ±<br>2.3<sup>a</sup><br>(n=5) | 39.2 ±<br>3.8 | 39.6 ±<br>3.4<sup>a</sup><br>(n=5) | 380.9 ±<br>70.2 | 391.1 ±<br>64.8<sup>a</sup><br>(n=5) | -58.2 ±<br>0.9 | -58.2 ±<br>1.4<sup>a</sup><br>(n=5) |
| Opto +<br>anti<br>S100β AB | 83.4 ±<br>3.2 | 70.0 ±<br>5.0<sup>a</sup><br>(n=4) | 35.3 ±<br>8.7 | 37.0 ±<br>4.8<sup>a</sup><br>(n=4) | 249.3 ±<br>58.3 | 204.7 ±<br>58.6<sup>a</sup><br>(n=4) | -56.5 ±<br>1.2 | -55.3 ±<br>1.1<sup>a</sup><br>(n=6) |
| Opto +<br>4.9 TTX | 74.3 ±<br>3.7 | 65.3 ±<br>7.6<sup>a</sup><br>(n=5) | 23.5 ±<br>3.7 | 21.4 ±<br>5.1<sup>a</sup><br>(n=5) | 233.3 ±<br>64.7 | 176.8 ±<br>49.3<sup>a</sup><br>(n=5) | -58.3 ±<br>0.8 | -58.2 ±<br>0.7<sup>b</sup><br>(n=5) |

### Antibody specificity

The specificity of the guinea-pig anti-S100β antibody 287004 was tested by the provider (Synaptic Systems) using immunocytochemistry and western blots on cells transfected with the S100β sequence and on brain slices. Its specificity was further confirmed using western blot analysis (Filice et al. 2017). The staining obtained in brain sections was reported to be similar to that obtained with another antibody directed against S100β raised in rabbit (Filice et al. 2017). The specificity of the goat anti-GFP antibody AB5450 was tested by the provider (Abcam) using immunoprecipitation and immunocytochemistry on transfected cells and on brain slices. Its specificity was further confirmed using immunohistochemistry (Pan et al. 2019). The specificity of the rabbit anti-NeuN antibody AB104225 was tested by the provider (Abcam) using immunocytochemistry on mouse brain sections. Its specificity was further confirmed using immunohistochemistry (Muñoz-Cabrera et al. 2019) and western blot analysis on mouse spinal cord tissue (Turner et al. 2014). The specificity of the rabbit anti-MAP2 antibody AB32454 was tested by the provider (Abcam) using immunocytochemistry, immunofluorescence and western blot analysis. Its specificity was further confirmed using immunohistochemistry analysis on primary mixed neuroglia cultures (Ceballos-Diaz et al. 2015). The specificity of the anti-S100β antibody S2532 was tested by the provider (Sigma) using ELISA and immunohistochemistry analyses. The specificity of the anti-Nav1.6 antibody ASC-009 was tested using immunostainings in mice (Tian et al. 2014). The specificity of the AB4674 against GFAP was tested by the provider (Abcam) using western blot and immunohistochemistry analyses. The specificity of the mouse anti-myosin heavy chain type IIA antibody SC-71 was confirmed by immunohistochemistry and immunoblot analysis studies (Schiaffino et al. 1989) and by the provider (Developmental Studies Hybridoma Bank) using immunoblots in several mammalian species.

### Electrical stimulation

Glass-coated tungsten microelectrodes (0.7-3.1 MΩ with 10-40 μm exposed tip) and a Grass S88 stimulator coupled to a Grass PSIU6 photoelectric isolation unit for controlling stimulation intensity (Astro Med) were used for electrical stimulation. The stimulation electrode was placed within layer 1 (L1) 100-200 µm lateral to the level of the recording site within L5, to elicit action potentials in L1 axons contacting the recorded L5PN. The electrical stimulation consisted of square pulses (0.2 ms duration) applied with a frequency of 1-50 Hz for 5 s. A pause of at least 1 min was given between two stimulation trains. The stimulation intensities ranged from 20 to 150 μA.

### Ca^2+^-sensitive electrodes

Electrodes were made of borosilicate glass pretreated with dimethylchlorosilane and dried at 120 °C for 2 h as previously described (Morquette et al. 2015). The tip of the pipette was filled with Ca^2+^ ionophore I cocktail A (Sigma-Aldrich). The rest of the electrode was filled with 2M of CaCl_2_. Before recording in the brain slice, the electrode was calibrated with aCSF solutions containing increasing concentrations of Ca^2+^ (0, 0.4, 0.8, 1.2, 1.6 and 2 mM) and the voltage for each concentration was recorded. The calibration curve was fitted with a logarithmic function. The electrodes used typically exhibited a potential jump higher than 20 mV between the two calibration solutions containing 0.2 or 2 mM of Ca^2+^ (Morquette et al. 2015). Measurements of [Ca^2+^]_e_ peaks were done using Clampfit (Molecular Devices).

### Electrophysiological data analysis

Recordings were analysed using standard scripts in Clampfit. For neurons, the passive properties included the resting membrane potential (in mV), the input resistance (in MΩ), and the membrane time constant Tau (in ms). The resting membrane potential was measured when no current was injected to the recorded neuron in current-clamp mode. The input resistance was determined as the voltage change induced by a small hyperpolarizing current (−20 to −40 pA) of 1 s duration applied from resting membrane potential divided by the amount of injected current. The membrane time constant was determined by fitting an exponential function to the first 200 ms of the voltage curve obtained in response to a small (−20 to −40 pA) hyperpolarizing current applied from resting membrane potential. Spikes were described from the current-voltage (IV) curve obtained in current clamp mode. The spike amplitude was measured using standard Clampfit script from the action potential evoked during a 1 s long depolarizing current pulse (20 to 100 pA). Spike amplitude was measured between spike peak and membrane potential value 5 ms before the peak.

### Statistics

Data in the text are presented as the mean ± standard error of the mean (sem). Correlations between variables and their significance as well as 95% confidence intervals were calculated using Sigma Plot 11.0. No statistical method was used to pre-determine sample sizes. The sample sizes in the present study are similar to those used in the field. No randomization or blinding procedure was used. Parametric analyses were used when assumptions for normality and equal variance were respected, otherwise non-parametric analyses were used. Two-tailed paired Student t-tests were performed for comparing means between two dependent groups. For more than two independent groups, a parametric one-way analysis of variance (ANOVA) or a non-parametric Kruskall-Wallis ANOVA were used. For more than two dependent groups, a parametric one-way analysis of variance (ANOVA) for repeated measures or a non-parametric Friedman ANOVA for repeated measures were used. When two factors were tested, a two-way ANOVA for repeated measures was performed. Both ANOVA analyses were followed by a Student-Newman-Keuls post-hoc test for multiple comparisons between groups. When group sizes were unequal, the Dunn’s test was used. Statistical differences were assumed to be significant when *P* < 0.05.

### Data availability

The datasets generated and analysed for the current study are available from the corresponding authors.

## Results

### Optogenetic stimulation of astrocytes induces depolarizations and increases in intracellular Ca^2+^ in astrocytes

To test the effects of astrocytic stimulation on neuronal excitability, we crossed homozygous mice expressing Cre recombinase under control of the GFAP promoter with homozygous mice expressing ChR2-EYFP flanked with a stop cassette and loxP sites. In the GFAP-ChR2-EYFP offspring, the stop cassette was excised by the Cre recombinase in cells expressing the GFAP promoter, and ChR2-EYFP was then expressed under control of the CAG promoter during cell lifetime. To determine whether the expression of ChR2-EYFP was restricted to astrocytes in GFAP-ChR2-EYFP mice, we performed double labelling experiments using immunohistochemistry against EYFP (see Methods) and against NeuN to label neuronal cell bodies in sections of V1. EYFP-positive processes formed a dense meshwork closely surrounding NeuN positive cell bodies that did not overlap with EYFP labeling in the vast majority of cases (n = 4 mice, Fig. 1A-F), but in rare cases faint EYFP labeling was seen overlapping a NeuN labeled neuron. When MAP2 was used to label neuronal processes (n = 4 mice); most neuronal fibers were negative to EYFP, but some rare areas of partial overlap were seen. We could not resolve whether these represented astrocytic EYFP positive processes running over neuronal parts or neurons positive to EYFP. However, to have a quantified estimation of such overlap, we combined NeuN, MAP2 and EYFP labeling in two mice and found that 94% of neuronal cell bodies were negative for EYFP (150/159 counted neurons). To further ensure that ChR2-EYFP did not accumulate in neuronal processes, but was located in astrocytes we carried out triple labelling immunohistochemistry experiments against MAP2, EYFP and S100β which labels mostly astrocytic cell bodies and proximal astrocytic processes (n = 2 mice). MAP2-positive neuronal processes (Fig. 1G,K) were surrounded by a dense EYFP-positive meshwork (Fig. 1H, L), but negative for EFYP (Fig. 1J, N) in contrast to putative astrocytes, immuno-positive to S100β (Fig. 1I, M), which were co-labelled with EYFP (Fig. 1J, N; n = 2 mice). To further confirm that ChR2-EYFP was not expressed in L5PN, we patched L5PN in slices and filled them with biocytin that was afterwards revealed and combined with immunostaining against EYFP. We found no patched L5PN positive for EYFP (n = 6 neurons from 4 mice; Fig. 1O-T). Altogether this indicated that ChR2-EYFP expression was specific to astrocytes in GFAP-ChR2-EYFP mice.

**Figure 1.**
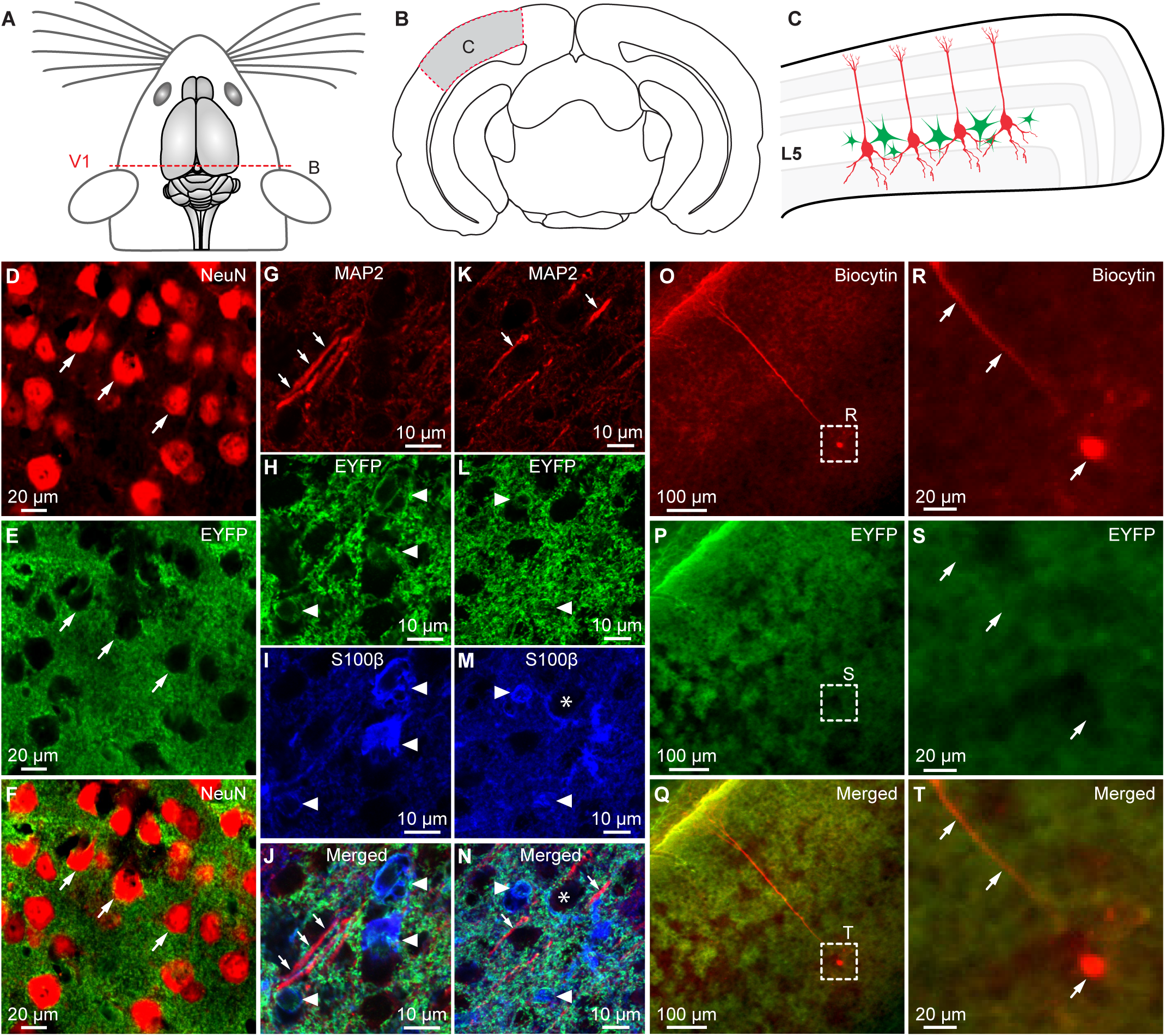
Selective expression of channelrhodopsin (ChR2) in astrocytes of GFAP-ChR2 mice. A. Dorsal view of the mouse brain showing the level at which the coronal slice of primary visual cortex (V1) was taken. **B.** Coronal brain slice, showing in the grey area enclosed by red dashed lines, the portion of V1 enlarged in C. **C.** Location of pyramidal neurons (red) and astrocytes (green) within layer 5 (L5). **D-F.** Immunofluorescence experiments against the enhanced yellow fluorescence protein (EYFP) that was fused with channelrhodopsin (ChR2) in GFAP-ChR2-EYFP mice. The arrows point to examples of neurons (NeuN-positive) that were negative for EYFP immunostaining. **G-J, K-N.** Two examples of triple labelling experiments in GFAP-ChR2 mice, showing that neuronal fibers positive for MAP2 (in red, see arrows in G, J, K, N) were negative for EYFP (in green, H, J, L, N) and negative for S100β (in blue, I, J M, N). S100β-positive cells were positive for EYFP (arrowheads in H-J, L-N). The asterisks in M-N delineates a putative neuron (i.e. negative for S100β and EYFP) closely surrounded by a glial process positive both for S100β and EYFP. J and N illustrate the merged three markers from G-I and K-M respectively. **O-T.** Double labelling experiments showing a typical L5PN filled with biocytin after streptavidin revelation at low (O-Q) and high (R-T) magnification in GFAP-ChR2 mice. Q and T illustrate the merged two markers from O-P and R-S respectively. The arrows in R-S point to the typical apical dendrite and soma of the recorded L5PN.

We then established, using whole-cell patch recordings from identified astrocytes (see Methods), that optogenetic stimulation (440-480 nm, see legend of Fig. 2) depolarized L5 astrocytes (7.3 ± 1.7 mV, n = 6 astrocytes from 6 slices from 5 mice; Fig. 2A-C) and induced Ca^2+^ responses in V1 slices loaded with Rhod-2-AM, a Ca^2+^ indicator preferentially taken up by astrocytes (Gordon et al. 2009; Mulligan and MacVicar 2004). Focal optogenetic (440-480 nm) stimulation of areas restricted to ∼20-30 µm in diameter to encompass only 2-3 astrocytes, evoked strong Ca^2+^ responses in several Rhod-2-positive cells over an area of ∼20-320 µm in diameter (Fig. 2D-E; n = 10 slices from 10 mice). Patch-clamp recordings in 3 of these Rhod-2-positive cells (from 3 slices, from 3 mice), ensured that they were astrocytes.

**Figure 2.**
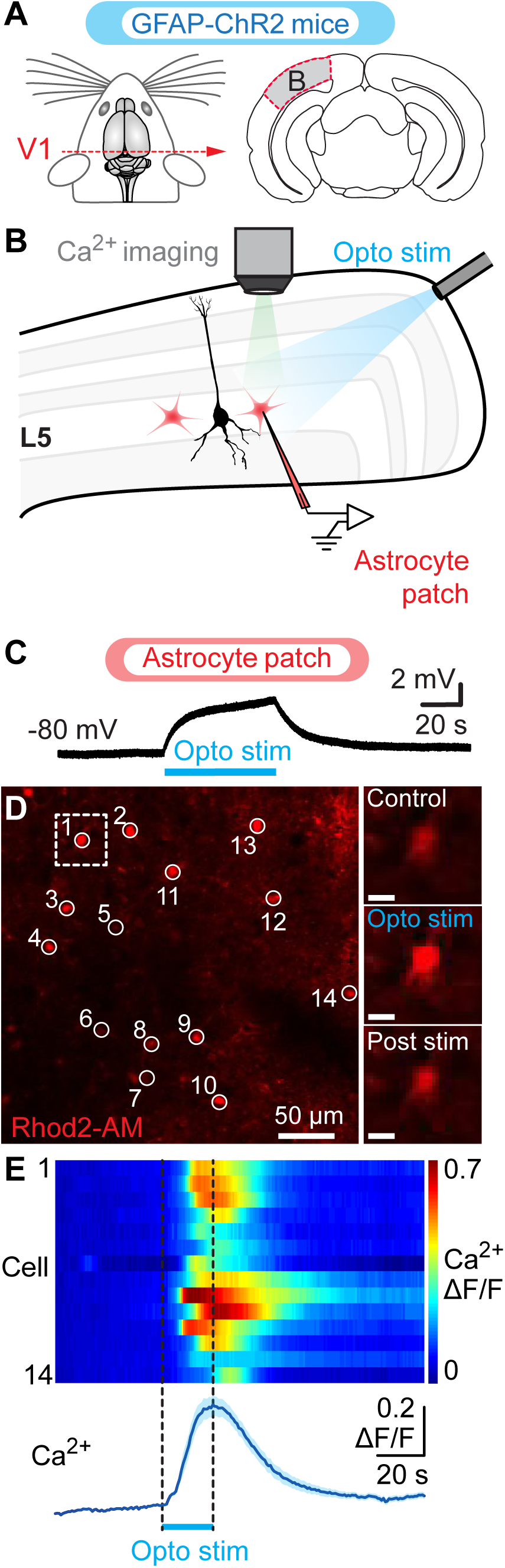
Optogenetic stimulation of astrocytes elicits calcium responses and depolarizes layer 5 astrocytes in GFAP-ChR2 mice. A-B. Schematic representation of the experimental set-up for calcium (Ca^2+^) imaging and whole-cell patch recordings from astrocytes (red cells) during optogenetic stimulation of astrocytes in GFAP-ChR2 mice (see Methods and Fig. 1). **C-D.** Blue light stimulation (440-480 nm, 20-60 s pulse, 10 % laser power) depolarized a patched astrocyte (C) and elicited Ca^2+^ responses (D) in ensembles of L5 cells (encircled in white) loaded with the Ca^2+^ indicator Rhod2-AM that is preferentially taken up by astrocytes (Gordon et al. 2009). Right of D, increase in Ca^2+^ fluorescence evoked in cell 1 by blue light (440-480 nm, 20 s pulse, 20 % laser power). **E.** Color plot illustrating the Ca^2+^ responses (ΔF/F) of Rhod2-AM-positive cells in response to astrocytic optogenetic activation with blue light (440-480 nm, 20 s pulse, 20 % laser power). Each line illustrates the response of individual cells numbered from top to bottom. Warmer colors (red) indicate larger Ca^2+^ responses. Bottom, average ± sem Ca^2+^ response for the 14 cells shown in D.

### Optogenetic activation of astrocytes elicits spiking in L5PN

L5PN were then targeted for patch-clamp recordings on the basis of the location of their soma and their dendritic morphology (Fig. 1K) to assess the effects of astrocytic stimulation on L5PN activity (Fig. 3A). Blue light pulses as short as 5 sec evoked a rapid (latency to first spike 0.42 ± 0.20 s, n = 6 neurons) firing response (Fig. 3B) that ranged from 0.4 to 141.9 Hz. In most cases, the evoked spiking occurred at a frequency below 40 Hz (80% of all interspike intervals (ISIs), 132/166 ISIs). The remaining 20% of the ISIs displayed higher frequencies (34/166 ISIs pooled from 6 trials obtained from the 6 neurons) with occasional bursts of action potentials (2 or more spikes with an interburst frequency of ∼1-40 Hz) where intraburst firing frequency reached more than 100 Hz. Such high frequency bursts have been reported in L5PN and are believed to play a role in coupling information originating from distinct cortical layers (e.g. Larkum et al. 1999, Schubert et al. 2001, Jacob et al. 2012).

**Figure 3.**
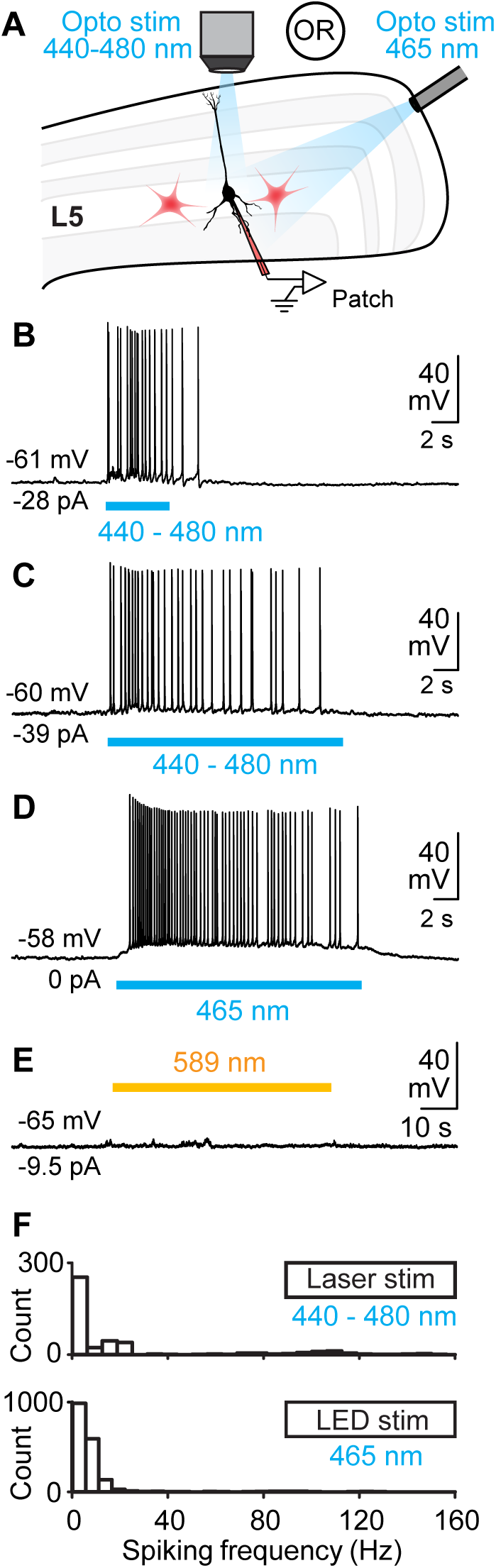
Optogenetic stimulation of astrocytes evoke spiking responses in patched layer 5 pyramidal neurons (L5PN) of GFAP-ChR2 mice. A. Scheme of the coronal brain slice, where the cortical layers in V1 are indicated. **B, C.** Tonic spiking activity elicited by astrocytic ChR2 activation with a short (5 s; B) and a long (20 s, C) pulse of laser stimulation (440-480 nm). **D.** Blue light stimulation with an LED (465 nm, 20 s pulse, 1000 mA LED intensity) evoked a similar response to laser stimulation. **E.** Absence of effect on L5PN electrophysiological activity when using a different wavelength (589 nm, 60 s pulse 10 % laser power) for light stimulation. Distribution of the instantaneous spiking frequencies of 55 spiking responses elicited in 55 L5PN when activating astrocytic ChR2 with either a laser (440-480 nm, 5 or 20 s pulses, 1-25 % laser power; upper graph) or a LED (465 nm, 20 s pulses, 400-1000 mA LED intensity; lower graph).

Then we tested longer stimulation pulses (20 s) to assess whether astrocytic stimulation affected inputs integration in L5PN over longer time periods, but first ensured that these elicited similar effects to shorter pulses. In 55 L5PN (from 54 slices from 41 GFAP-ChR2 mice), which electrophysiological properties are summarized in Table 1, blue light stimulation with a laser focused on smaller areas (Fig. 3C) or with a 50 μm optic fiber (Fig. 3D) yielded L5PN depolarizations and robust firing similar to those observed with shorter pulses with no signs of neuronal degradation with time. Photostimulation of large areas with other light wavelengths (589 nm, 60 s pulse or 559 nm, 20 s pulse, 10 % laser power in both cases) failed to elicit any electrophysiological response in L5PN (n = 2 neurons from 2 slices from 2 mice, Fig. 3E). As with shorter light pulses, firing frequencies of the elicited responses ranged from 0.1-156.5 Hz, and the evoked spiking occurred mostly at a frequency below 40 Hz (Fig. 3F). Higher frequency spiking accounted for 5% of all interspike intervals (ISIs; n = 114/2244 ISIs pooled from 55 trials obtained from 55 neurons). The first spike appeared on average 1.42 ± 0.23 s (n = 55 neurons) after the beginning of the blue light pulse. Altogether these results establish that spiking in L5PN were evoked only when astrocytes were illuminated with 440-480 nm light. This activated ChR2, an opsin that was specifically expressed in astrocytes in our GFAP-ChR2-EYFP mice as confirmed by our anatomical experiments above.

**Table 1.** Neuronal properties for the three types of mouse strains. R, resistance, Tau, membrane time constant, Vm, membrane potential. **^a^**P < 0.05 vs. GFAP-ChR2, Dunn’s test either after a Kruskal-Wallis one-way ANOVA on ranks (*P* < 0.05). **^b^**P < 0.05 vs. Thy1-GCaMP6f, Dunn’s test either after a Kruskal-Wallis one-way ANOVA on ranks (*P* < 0.05). **^c^**P < 0.05 vs. GFAP-ChR2, Dunn’s test either after a Kruskal-Wallis one-way ANOVA on ranks (*P* < 0.001). **^d^**P < 0.05 vs. Thy1-GCaMP6f, Dunn’s test either after a Kruskal-Wallis one-way ANOVA on ranks (*P* < 0.001). **^e^**P < 0.01 vs. GFAP-ChR2, Student-Newman-Keuls test after a one-way ANOVA (*P* < 0.001).

|  | N<br>cells | Age<br>(days) | R<br>(MΩ) | Spike<br>Amplitude<br>(mV) | Tau<br>(ms) | Vm<br>(mV) |
| --- | --- | --- | --- | --- | --- | --- |
| **GFAP**<br>**ChR2** | 70 | P19.8 ±<br>0.6 | 260.0 ±<br>16.0 | 81.2 ± 1.2 | 38.1 ±<br>1.8 | -58.1 ±<br>0.4 |
| **Thy1-**<br>**GCaMP6**<br>**f** | 20 | P20.0 ±<br>1.1 | 232.8 ±<br>25.6 | 87.4 ± 1.7<sup>a</sup> | 39.6 ±<br>3.4 | -61.0 ±<br>1.1<sup>e</sup> |
| **WT** | 21 | P17.5 ±<br>0.5 | 316.2 ±<br>34.5 | 81.0 ± 2.1<sup>b</sup> | 54.3 ±<br>3.7<sup>c,d</sup> | -61.0 ±<br>0.7<sup>e</sup> |

### Optogenetic activation of astrocytes potentiates L1 inputs in L5PN

We then tested whether activation of astrocytes located in the proximal L5 region influenced integration of distal inputs to L5PN (Fig. 4A-B) elicited with electrical stimulation with an electrode placed in L1 slightly lateral (∼100 µm) to the apical tuft of the recorded neuron (Fig. 4A). Stimulation intensity in L1 was set to obtain ∼20% success rate (i.e. number of time a stimulus pulse delivered in L1 elicited a somatic spike within a 50 ms time window afterwards divided by the number of L1 stimulation pulses × 100; Fig. 4B, C, I), while laser intensity for optogenetic stimulation of astrocytes was carefully reduced to elicit a subthreshold depolarization that could be detected but that was as much as possible insufficient to elicit spiking by itself. This caused a 1.3 ± 0.3 mV depolarization measured in 7 L5PNs in from 6 slices from 6 mice (control: −59.8 ± 0.8; opto: −58.5 ± 0.8, post opto: −60.0 ± 0.9 mV, *P* < 0.05 for control vs. opto stim and for opto stim vs. post stim, Student-Newman-Keuls test after a one-way ANOVA for repeated measures, *P* < 0.001) and only 1.3 ± 1.1 spikes. When applied together with L1 stimulation, such low optogenetic activation of astrocytes increased the success rate of L1-evoked synaptic responses from 16.3 ± 4.7 to 35.3 ± 8.8 % (*P* < 0.05, Student-Newman-Keuls test after a one-way ANOVA for repeated measures, *P* < 0.05, n = 10 neurons from 8 slices from 8 mice) (Fig. 4B, D, I). The potentiating effect of astrocytic stimulation lasted as long as the duration of the light pulse, and success rate of L1 stimulation returned to lower values when the light was switched off (9.0 ± 5.3 %; *P* < 0.05 vs. optogenetic stimulation; Fig. 4B, E, I). This indicated that astrocyte activation increased the ability of distal inputs to elicit spikes at the soma by depolarizing the soma.

**Figure 4.**
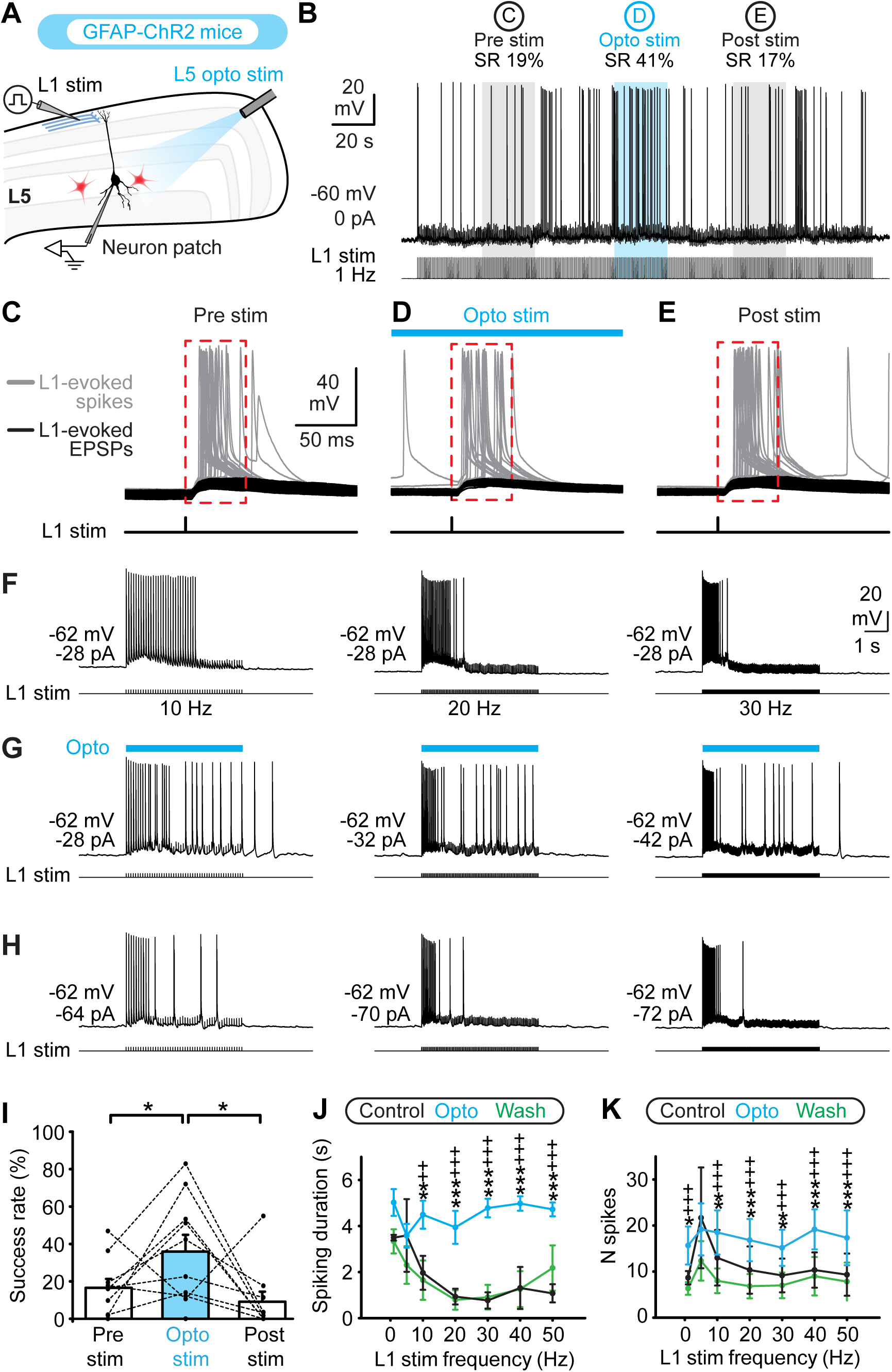
Optogenetic stimulation of layer 5 astrocytes potentiates inputs of layer 1 (L1) to layer 5 pyramidal neurons (L5PN). A-B. In coronal slices of the primary visual cortex of GFAP-ChR2 mice, a tungsten electrode was placed in L1 to stimulate L1 axons (1 Hz trains, 0.2 ms pulses, 20-150 µA) contacting the recorded L5PN. The stimulation intensity was set to have a low success rate (B, left). Most L1 pulses induced EPSPs (black traces in C, D, and E), and spikes (grey traces) were elicited in about 20% of trials. **C.** Because an increase in spiking activity was expected to occur during optogenetic stimulation, we quantified the success rate of spikes evoked by L1 stimulation that appeared in a 50 ms time window (red dashed rectangle in C, D, and E) after each single pulse delivered to L1. Activation of ChR2 in astrocytes was induced with blue light (440-480 nm, 20-95 s pulses, 5-25 % laser power) applied locally around the cell body of the recorded L5PN. The success rate was measured during 21 to 137 s in control conditions (B, C), 20 to 95 s in optogenetic stimulation condition (B, D), and during 10 to 157 s after blue light was switched off (B, E). **F-H.** Firing evoked by electrical stimulation of L1 axons (at 10, 20, 30 Hz, 0.2 ms pulses, 5 s train, 70-100 µA) contacting the recorded L5PN were investigated in control condition (F), during optogenetic activation of astrocytes (440-480 nm, 5 s pulse, 1-12% laser power) near the recorded L5PN (G), and after blue light was switched off (H). **I.** Effect of optogenetic activation of astrocytes on the success rate of L1 stimulation in the pooled data (n = 10 neurons for 8 slices from 8 mice, \**P* < 0.05, Student-Newman-Keuls test after a one-way ANOVA for repeated measures, *P* < 0.001). **J.** The duration of the spiking response was compared for each L1 stimulation frequency (\*\**P* < 0.01, \*\*\**P* < 0.001 vs. control; ^++^*P* < 0.01, ^+++^*P* < 0.001 vs. drug condition, Student-Newman-Keuls test after a two-way ANOVA for repeated measures on ranks *P* < 0.001). **K.** The number of spikes were compared for each L1 stimulation frequency (\**P* < 0.05, \*\**P* < 0.01, \*\*\**P* < 0.001 vs. control; ^+++^*P* < 0.001 vs. optogenetic stimulation, Student-Newman-Keuls test after a two-way ANOVA for repeated measures on ranks *P* < 0.001). In J-K, data were pooled from 6 neurons from 4 slices from 3 mice.

### Optogenetic activation of astrocytes overrides the adaptation of firing evoked by L1 stimulation

To assess how astrocytic activation affects neuronal responsiveness over time, we examined if it altered the responses of L5PN to repetitive stimulation of L1 inputs. This time, the intensity of L1 stimulation was adjusted to generate AP in L5PN (Fig. 4F) and responses to 5 s trains of stimuli delivered at different frequencies (1 to 50 Hz) were recorded. At 1, 5 and 10 Hz, L5PN generated a spike or a doublet at nearly every stimulus pulse, but for higher stimulation frequencies (20, 30, 40 and 50 Hz), their discharge adapted, and the number of spikes rapidly decreased or firing completely ceased during the stimulus train (Fig. 4F). Optogenetic activation of astrocytes near the soma of the recorded L5PN overrode the firing adaptation at higher frequencies (Fig. 4G) as shown by measures of spiking response duration (Fig. 4J, for 10, 20, 30, 40, and 50 Hz, *P* < 0.01 to *P* < 0.001 vs. control, Student-Newman-Keuls test after a two-way ANOVA for repeated measures on ranks, *P* < 0.001) and measures of the number of spikes (Fig. 4K, for 1, 10, 20, 30, 40, and 50 Hz, *P* < 0.05 to *P* < 0.001 vs. control, Student-Newman-Keuls test after a two-way ANOVA for repeated measures on ranks, *P* < 0.01, n = 6 neurons from 4 slices from 3 mice). These effects disappeared after blue light was switched off (Fig. 4H) both for the spiking response duration (Fig. 4J, for 10, 20, 30, 40, 50 Hz, *P* < 0.01 to *P* < 0.001 vs. optogenetic stimulation) and for the number of spikes (Fig. 4K; for 1, 10, 20, 30, 40, and 50 Hz, *P* < 0.001 vs. optogenetic stimulation). This suggests that astrocyte activation proximal to the soma of L5PN prolong the time window over which they can integrate distal inputs.

### Astrocyte-induced spiking in L5PN does not depend solely on glutamatergic, GABAergic or purinergic signaling

Next, we examined how astrocytic activation produced the above-described effects. Astrocytes can influence neural activity through release of different gliotransmitters that include glutamate (Bezzi et al. 1998; Parpura et al. 1994; Poskanzer and Yuste 2011), GABA (Kozlov et al. 2006; Liu et al. 2000), ATP (Newman 2003; Panatier et al. 2006; Perea et al. 2014; Poskanzer and Yuste 2011) or the NMDA co-agonist D-Serine (Henneberger et al. 2010; Yang et al. 2003; for review, see Araque et al. 2014). At least two of these gliotransmitters, glutamate and ATP, have been shown to be released following optogenetic manipulations of astrocytes (Berlinguer-Palmini et al. 2014; Chen et al. 2013a; Gourine et al. 2010; Gradinaru et al. 2009; Li et al. 2012; Perea et al. 2014; Poskanzer and Yuste 2016; Sasaki et al. 2012; Tan et al. 2017) and a recent study showed that neuronal excitation following light stimulation of hippocampal astrocytes expressing ChR2 resulted from a synergistic action of purinergic signalling on glutamate release (Shen et al. 2017).

Thus, we started by testing involvement of fast glutamatergic and GABAergic transmissions with bath applications of CNQX (10 µM), AP5 (26 µM), and gabazine (20 µM) that block AMPA/kaïnate, NMDA and GABA_A_ receptors respectively. Consistent with previous report that astrocytic glutamate release can activate nearby cortical neurons (e.g. Poskanzer and Yuste 2011), we observed a decrease in the number of spikes evoked by optogenetic astrocyte activation in presence of the blockers (control 37.3 ± 10.2 vs. drugs 18.8 ± 7.4 spikes, *P* < 0.05, paired t-test, n = 7 neurons from 7 slices from 7 mice, 1 to 4 trials per drug condition, Fig. 5A-C). We observed no significant modification of input resistance, membrane time constant (Tau) or of spike amplitude following the application of blockers (*P* > 0.05 vs. control, paired t-test or Wilcoxon signed rand test, n = 5 neurons tested out of 7, Table 2). However, since some firing persisted in the presence of blockers, and given the fact that effects of glutamate released from astrocytes are often mediated by metabotropic glutamate receptors (mGluRs) (Perea et al. 2014), we tested another cocktail that contained a wide spectrum mGluR antagonist LY 341495 (100 µM) and the other blockers (CNQX 10 µM, AP5 50 µM, LY 341495 100 µM, gabazine 20 µM). Unexpectedly, the number of spikes elicited in L5PN by optogenetic activation of astrocytes did not change significantly even after a 30 min bath application of these blockers (control 38.2 ± 11.9 vs. 46.5 ± 24.0 spikes, *P* > 0.05, paired t-test, n = 4 slices from 4 mice, 1 to 3 trials per drug condition, Fig. 5D-F), nor did the input resistance, membrane Tau or spike amplitude (*P* > 0.05 vs. control, paired t-test or Wilcoxon signed rank test, n = 4 neurons tested out of 4, Table 2), suggesting that metabotropic and ionotropic receptors are producing opposing effects on L5PN output by perhaps acting on different sets of cells or causing opposite effects on the cells. However, in all cases, some firing evoked by astrocyte optogenetic activation persisted after blockade of these receptors.

**Figure 5.**
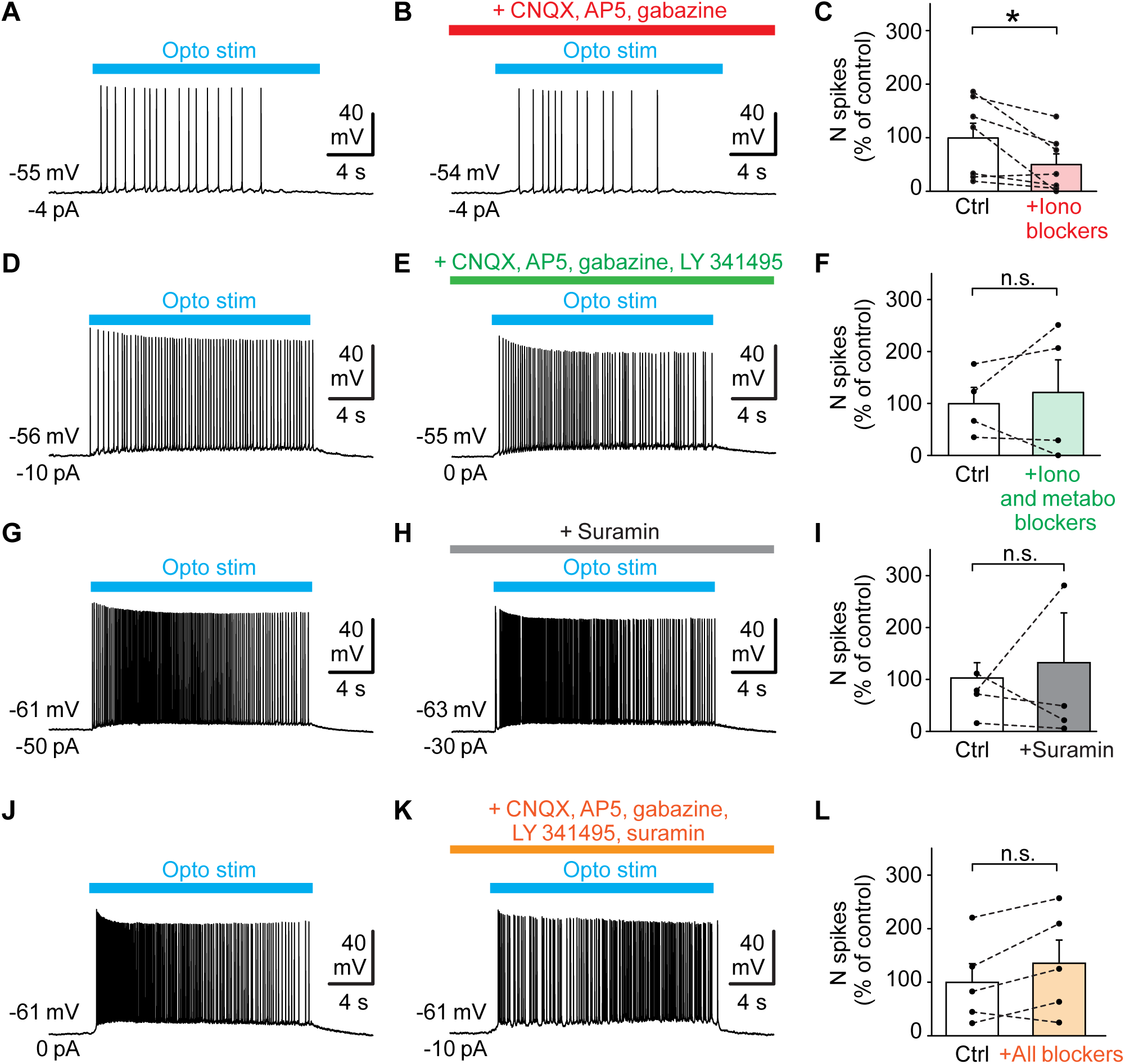
Spiking evoked in layer 5 pyramidal neurons (L5PN) by optogenetic activation of L5 astrocytes persists in presence of glutamatergic, GABAergic and purinergic blockers. A-C. Bath application (30 min) of a cocktail containing the AMPA/kaïnate receptor antagonist CNQX (10 μM), the NMDA receptor antagonist AP5 (26 μM) and the GABA_A_ receptor antagonist gabazine (20 µM) reduced the number of spikes evoked by optogenetic stimulation of astrocytes with blue light (465 nm, 20 s pulses, 800 mA LED intensity) in slices of GFAP-ChR2 mice (* *P* < 0.05, paired t test, n = 7 neurons) but did not abolish spiking activity. **D-F.** Bath application (30 min) of a cocktail containing the AMPA/kaïnate receptor antagonist CNQX (10 μM), the NMDA receptor antagonist AP5 (50 μM), the GABA_A_ receptor antagonist gabazine (20 µM) and the metabotropic receptor blocker LY 341495 (100 µM) did not abolish the spiking activity (n.s. *P* > 0.05, paired t test, n = 4 neurons). **G-I.** Bath application (30 min) of the purinergic receptor inhibitor suramin (100 µM) did not abolish the spiking activity evoked by optogenetic stimulation of astrocytes with blue light (465 nm, 20 s pulses, 800 mA LED intensity) (n.s. *P* > 0.05, paired t test, n = 4 neurons). **J-L.** Bath application (30 min) of a cocktail containing the AMPA/kaïnate receptor antagonist CNQX (10 μM), the NMDA receptor antagonist AP5 (50 μM), the GABA_A_ receptor antagonist gabazine (20 µM), the metabotropic receptor blocker LY 341495 (100 µM) and the purinergic blocker suramin (100 µM) did not abolish the spiking activity (n.s. *P* > 0.05, paired t test, n = 5 neurons). Data from A-B, D-E, G-H and J-K are from 4 different slices from 4 different mice.

We then tested for involvement of ATP release with bath application (30 min) of the wide-spectrum purinergic receptor blocker, suramin (100 µM), which did not significantly change the number of spikes evoked by optogenetic activation of astrocytes (41.2 ± 11.8 vs. suramin 52.9 ± 38.3 spikes, *P* > 0.05, paired t-test, n = 4 neurons from 4 slices from 4 mice, 1 to 3 trials per drug condition Fig. 5G-I). No significant modification of input resistance, membrane Tau or spike amplitude was observed (*P* > 0.05 vs. control, paired t-test or Wilcoxon signed rank test, n = 4 neurons tested out of 4, Table 2). Finally, to insure that neither glutamate, nor GABA, nor ATP release were involved, we bath applied (30 min) all five blockers together (CNQX 10 µM, AP5 50 µM, LY 341495 100 µM, gabazine 20 µM, suramin 100 µM) and found no modification in the number of spikes evoked by optogenetic activation of astrocytes (56.9 ± 19.9 vs. 77.2 ± 24.7 spikes, *P* > 0.05, paired t-test, n = 5 slices from 5 mice, 1 to 3 trials per condition, Fig. 5J-L). Again, we observed no significant modification of input resistance, membrane Tau or spike amplitude (*P* > 0.05 vs. control, paired t-tests, n = 5 neurons tested out of 5, Table 2).

Thus, at most, we observed a ∼50% decrease in the number of spikes evoked by astrocyte optogenetic activation in presence of blockers of fast glutamatergic and GABAergic transmission, which is in accordance with previous reports showing that astrocytes can modify neuronal activity with these transmitters (Bezzi et al. 1998; Kozlov et al. 2006; Liu et al. 2000; Parpura et al. 1994; Poskanzer and Yuste 2011), but no complete block was observed, suggesting that under the conditions used, an additional mechanism could contribute to the observed firing in L5PN following optogenetic stimulation of astrocytes.

### Spiking evoked in L5PN by optogenetic activation of astrocytes depends on S100β and Nav1.6 channels

We have previously shown, in a brainstem sensorimotor circuit, that astrocytes can modulate neuronal firing by potentiating a sodium persistent current (I_NaP_) through release of S100β (Morquette et al. 2015). S100β is an astrocytic calcium-binding protein which decreases the extracellular calcium concentration ([Ca^2+^]_e_), when released in the extracellular space, and by doing so, activates I_NaP_. To determine whether [Ca^2+^]_e_ changes were involved in the observed effect, we placed a Ca^2+^-sensitive electrode near the recorded L5PN and recorded [Ca^2+^]_e_ during spiking episodes elicited by optogenetic stimulation of astrocytes in GFAP-ChR2-EYFP mice (Fig. 6A). Indeed, optogenetic activation of astrocytes decreased the [Ca^2+^]_e_ from 1.24 ± 0.02 to 1.11 ± 0.02 mM (*P* < 0.001 vs. control, Student-Newman-Keuls test after a one way ANOVA for repeated measures, *P* < 0.001, n = 6 slices from 5 mice) (Fig. 6A-C). [Ca^2+^]_e_ increased back to 1.26 ± 0.02 mM, 20 s after the end of the blue light pulse (*P* < 0.001 vs. optogenetic stimulation, n = 6 slices from 5 mice; Fig. 6A-C). The decrease in [Ca^2+^]_e_ was associated in time with the occurrence of the spiking activity in neighboring L5PN (Fig. 6B-C, n = 6 neurons, from 6 slices from 5 mice) and corresponded almost exactly to [Ca^2+^]_e_ decreases (delta decrease of 0.13 ± 0.02 mM) observed in the extracellular space of layer 2/3 of the somatosensory cortex of mice transitioning from sleep to awake states (Ding et al. 2016).

**Figure 6.**
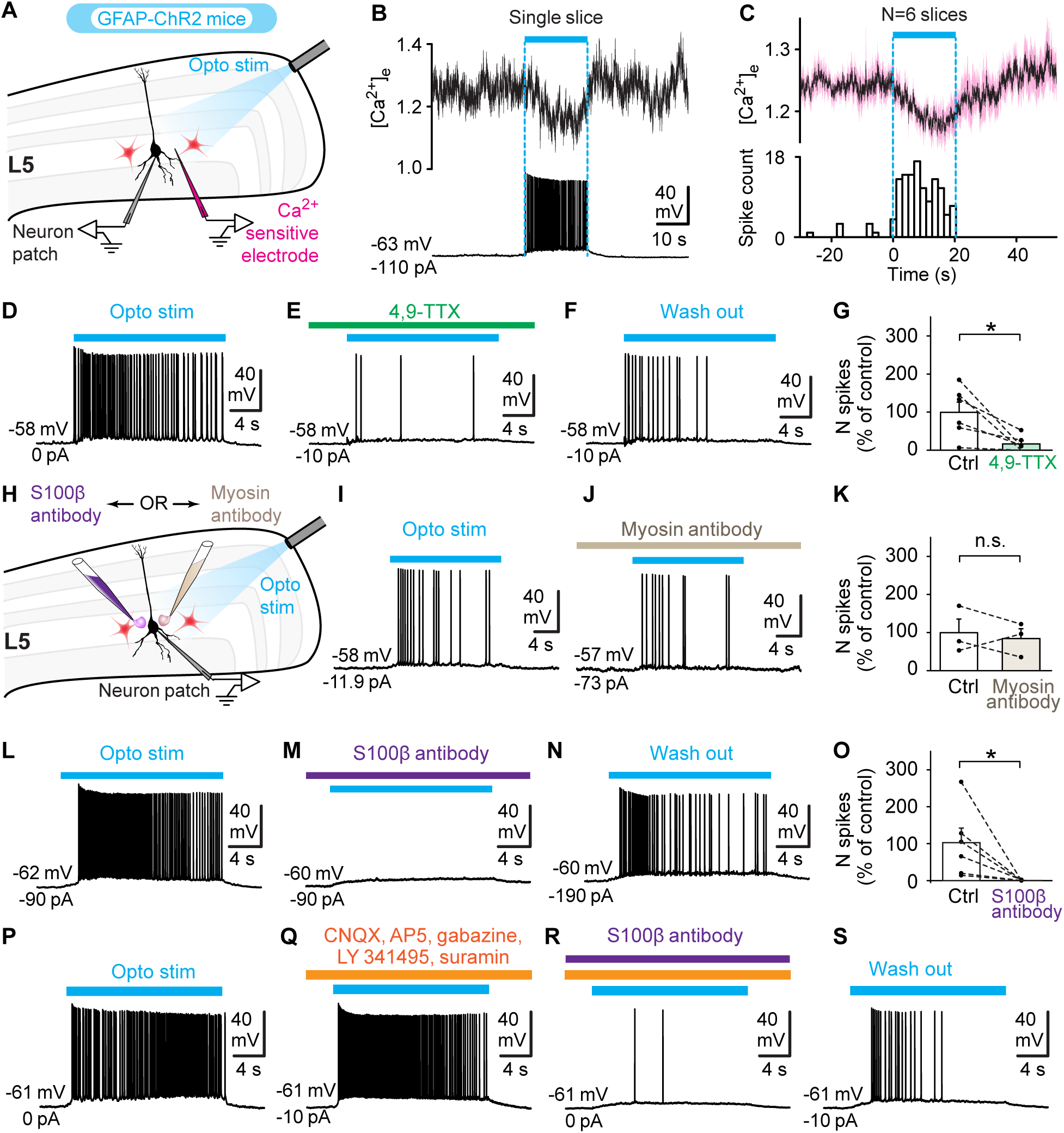
Optogenetic stimulation of layer 5 astrocytes decreases [Ca^2+^]_e_ and elicits S100β- and Nav1.6-dependent spiking in layer 5 pyramidal neurons (L5PN). A. Schema showing position of the recording electrodes for L5PN and measures of [Ca^2+^]_e_ during L5 optogenetic stimulation. **B.** Single slice data showing that optogenetic stimulation of L5 astrocytes with blue light (465 nm, 400 mA, 20 s pulse) evoked a decrease in [Ca^2+^]_e_ and simultaneous spiking in a nearby L5PN. **C.** Data pooled from 6 slices showing the average decrease in [Ca^2+^]_e_ (mean in black ± sem in pink) together with a binned raster plot counting the mean number of spikes evoked in 6 L5PN with blue light (465 nm, 400-800 mA, 20 s pulse). **D-G.** Spiking elicited in a L5PN by optogenetic stimulation (465 nm, 20 s pulses, 800 mA LED intensity) of L5 astrocytes is nearly abolished under 4,9-TTX (0.1 µM, 24 min) (E) and recovers after partial washout (F). In G, number (N) of optogenetically evoked (440-480 nm, 10% laser power, or 465 nm, 800 mA LED intensity, 20 s pulses) spikes in L5PN (\**P* < 0.05, paired t test, n = 7 neurons). **H.** Schema showing the experimental setup for local application of an anti-S100β antibody (40 μg/mL) or an unrelated anti-myosin antibody (477 μg/mL) during optogenetic stimulation. **I-K.** Local application of an anti-myosin antibody (20 min pulse, 0.1 psi) has no effect on the spiking elicited in an L5PN by optogenetic stimulation (440-480 nm, 20 s pulses, 10 % laser power) of astrocytes (n.s. *P* > 0.05, paired t test, n = 3 neurons). In K, n.s. *P* > 0.05, paired t test, n = 3 neurons. **L-O.** Spiking induced by stimulation of astrocytes (465 nm, 20 s pulses, 800 mA LED intensity) is abolished after local application of the anti-S100β antibody (8 min pulse, 0.1 psi) (M) and partially recovers after wash out (N). In O, \**P* < 0.05, paired t test, n = 6 neurons. **P-S.** Spiking induced by stimulation of astrocytes (465 nm, 20 s pulses, 800 mA LED intensity) persists in presence of glutamatergic, GABAergic and purinergic blockers (Q) but is abolished by local application of the anti-S100β antibody (30 min pulse, 0.1 psi) (R) and partially recovers after wash out of the drugs (S). Data from B, D-F, I-J and L-N and P-S were obtained from 4 different slices from 4 mice.

Several studies have shown that extracellular Ca^2+^ reduces Na^+^-single-channel current amplitude in a fast and voltage-dependent manner (Armstrong 1999; Armstrong and Cota 1991, 1999; Bahia et al. 2012; Corry 2013; Corry and Thomas 2012; Horn 1999; Tikhonov and Zhorov 2007; Vandenberg and Bezanilla 1991). This was shown to be of particular importance for I_NaP_-mediated firing (Brocard et al. 2013, 2016; Li and Hatton 1996; Morquette et al. 2015; Su et al. 2001). To determine whether astrocyte-evoked firing relied on I_NaP_, we tested its sensitivity to 4,9-anhydro-TTX (4,9-TTX), a specific blocker of sodium channels containing the Nav1.6 α subunit, because these are the channels most commonly associated to persistent Na^+^-currents (Chatelier et al. 2010). Bath application (7 to 50 min) of 4,9-TTX (0.1 µM) reduced the number of spikes elicited by optogenetic stimulation of astrocytes from 38.1 ± 10.2 to 6.5 ± 3.1 spikes (n = 6 neurons from 6 slices from 6 mice, 2 to 4 trials per drug condition, *P* < 0.05 vs. control, paired t test; Fig. 6D-G). No significant alteration of the input resistance, membrane Tau or of spike amplitude was detected in presence of 4,9-TTX (*P* > 0.05 vs. control, paired t-tests, n = 5 neurons tested out of 6, Table 2). The number of spikes partially recovered to 13.5 ± 5.9 spikes after washout of 4,9-TTX (n = 4 neurons from 4 mice, Fig. 6F).

We then tested whether S100β was involved in the astrocytic effect by applying an antibody directed against it to prevent its extracellular action (Condamine et al. 2018; Morquette et al. 2015; Sakatani et al. 2008). We have already shown, using isothermal titration calorimetry, that this antibody interferes with the ability of S100β to bind Ca^2+^ (Morquette et al. 2015). Local application of the anti-S100β antibody (40 μg/mL, 8-15 min pulse, 0.1 psi) dramatically decreased the number of spikes evoked in L5PN by optogenetic stimulation of astrocytes from 41.9 ± 16.0 to 0.2 ± 0.2 spikes (n = 6/6 neurons from 6 slices of 6 mice, 1 to 3 trials per drug condition, *P* < 0.05, paired t-test, Fig. 6H, L-O). This effect could be partially reversed in 2/2 neurons from 2 slices from 1 mouse (33.5 ± 12.5 spikes after wash out, Fig. 6N). No significant alteration of input resistance, membrane Tau or spike amplitude was detected in the presence of the anti-S100β antibody, Table 2). In 3 additional experiments, the spiking activity evoked by optogenetic activation of astrocytes that persisted in presence of bath-applied blockers of AMPA, NMDA, GABA_A_, mGluRs and purinergic receptors (CNQX 10 µM, AP5 50 µM, gabazine 20 µM, LY 341495 100 µM and suramin 100 µM) was dramatically decreased by local application of the anti-S100β antibody (65.3 ± 40.8 vs. 10.7 ± 9.7 spikes, 3 neurons from 3 slices of 3 mice; 1 to 3 trials per drug condition, Fig. 6P-R). Finally, to test the specificity of the effect of the anti-S100β antibody, we locally applied an antibody directed against myosin and found no effect on the spiking activity evoked by optogenetic stimulation of astrocytes (31.2 ± 11.1 vs 26.3 ± 8.0 spikes, n = 3 neurons from 3 slices of 2 mice; 1 to 3 trials per drug condition, Fig. 6H-K). Altogether these experiments demonstrate a key role for endogenous S100β and Nav1.6 in L5PN firing evoked by optogenetic activation of astrocytes.

### The astrocytic Ca^2+^-binding protein S100β and the calcium chelator BAPTA both decrease extracellular Ca^2+^ and evoke firing in V1 L5PN

We then tested whether small extracellular applications of the Ca^2+^-binding protein S100β or the Ca^2+^ chelator BAPTA, which both produce presumably small localised decreases of [Ca^2+^]_e_, are sufficient to induce activity in L5PN and to reproduce the effect of astrocyte optogenetic activation (Fig. 7A-B). Local applications of exogenous S100β (129 µM, 1 to 15 s pulses, 1 to 20 psi) within L5 elicited spiking in all 35 L5PN recorded (n = 16 neurons from 7 wild type mice, n = 16 neurons from 13 Thy1-GCaMP6f mice, and n = 3 neurons from 2 GFAP-ChR2 mice) (Fig 7C, E). Likewise, local applications of BAPTA within L5 (5-10 mM, 2-10 psi, 0.7 to 8 s pulses) elicited spiking in all 17 L5PN recorded (n = 2 neurons from 2 wild type mice; n = 11 neurons from 9 Thy1-GCaMP6f mice and n = 4 neurons from 3 GFAP-ChR2 mice; Fig. 7D, F). As was the case with optogenetic stimulation of astrocytes (Fig. 3), tonic spiking constituted most of the responses elicited by S100β and BAPTA with frequencies in the same range (0.3-134.4 Hz for S100β, 0.5-105.8 Hz for BAPTA, vs. 0.1-156.5 Hz for astrocyte optogenetic stimulation; Fig 7E, F). Higher firing frequencies were observed in doublets or burst-like firing in only a minor proportion of trials (159/980 ISIs, 16% of all ISIs, recorded in 35 trials from n = 35 neurons with S100β, and 19/1029 ISIs recorded in 17 trials from n = 17 neurons with BAPTA). Because S100β-evoked and BAPTA-evoked spiking activities were observed in the three mouse strains (Fig. 7E, F), and because electrophysiological properties of L5PN were largely similar in the three strains (Table 1), data from all three strains of mice were pooled in subsequent experiments.

**Figure 7.**
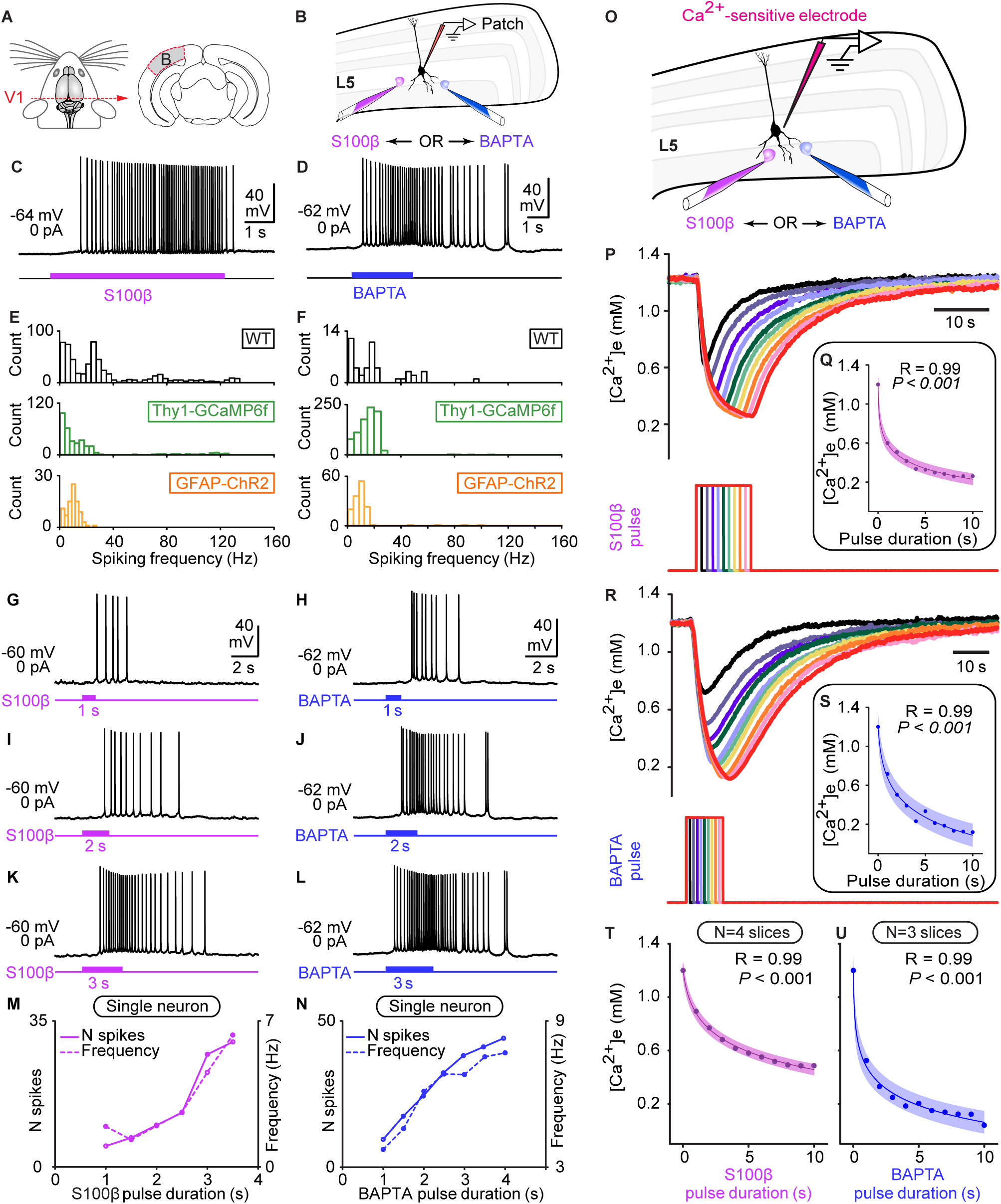
The astrocytic protein S100β elicits spiking activity in V1 layer 5 pyramidal neurons (L5PN) that is proportional to its effect on extracellular concentration of calcium [Ca^2+^]_e_. A. Drawing of the level at which sections were taken from the mouse brain to obtain slices of V1. **B.** Schema of the positions of the recording electrode and of the pipettes for local applications of the astrocytic protein S100β (purple) or the Ca^2+^ calcium chelator BAPTA (blue) near L5PN (black). **C, D.** Tonic spiking activity elicited by local application of S100β (C, 129 μM, 6 s pulse, 5 psi) or BAPTA (D, 5 mM, 3 s pulse, 5 psi). **E, F.** Distribution of the instantaneous spiking frequencies elicited by S100β (E) or BAPTA (F). **G-L.** Increase in S100β (129 µM, 5 psi, G, I, K) or BAPTA (5 mM, 5 psi, H, J, L) pulse duration increases the number of spikes and spiking frequency in L5PN. Data are from 2 different slices. **M-N.** Plot illustrating the number of spikes (solid line) or the spiking frequency (dashed line) as a function of S100β pulse duration (M) or BAPTA pulse duration (N) for the raw data illustrated respectively in G, I, K and H, J, L. **O.** Scheme of the coronal brain slice, where the cortical layers in V1 are indicated. An ion-sensitive electrode was placed within layer 5 (L5) to measure the variations of [Ca^2+^]_e_ elicited by local application of the astrocytic protein S100β or the Ca^2+^ chelator BAPTA. **P, R.** Variations in [Ca^2+^]_e_ evoked by applications of S100β (P; 129 μM, 10 psi) or BAPTA (R; 5 mM, 5 psi) of increasing durations (1 to 10 s). For each pulse duration tested, the corresponding signal in [Ca^2+^]_e_ appears with the same color. **Q, S.** Relationship between the duration of S100β (Q) or BAPTA (S) pulse and the [Ca^2+^]_e_ in the data shown in P and R respectively. **T-U.** Relationship between the average duration of S100β (T) or BAPTA (U) pulse and the average [Ca^2+^]_e_ measured in 4 and 3 slices respectively. In Q, S, T, U the fit and the 95% prediction intervals are shown.

### S100β exerts a precise control of [Ca^2+^]_e_ and of L5PN spiking activity in V1

The fact that S100β and BAPTA elicited similar effects on L5PN spiking activity suggests that these effects resulted from their Ca^2+^ binding ability. Thus, we tested how applications of variable duration of both drugs affected [Ca^2+^]_e_ (Fig. 7O) and whether such changes were reflected in the elicited firing of L5PN (Fig. 7G-N). Local applications of increasing duration (10 pulses of 1-10 s) of either S100β (129 µM, 10 psi) or BAPTA (5 mM, 5 psi) produced larger and longer lasting decreases of [Ca^2+^]_e_ (Fig. 7P, R). In single cases (Fig. 7Q,S), as in pooled data (Fig. 7T-U), the relationship between [Ca^2+^]_e_ and the duration of local applications of either drugs followed a logarithmic function (for the pooled data, R = 0.99, *P* < 0.001 for S100β, n = 4 slices from 4 mice, and R = 0.99, *P* < 0.001 for BAPTA, n = 3 slices from 3 mice). At the concentration used, S100β (129 µM) was slightly less potent than BAPTA (5 mM) and decreased [Ca^2+^]_e_ from 1.2 to approximately 0.5 mM, whereas BAPTA could decrease it further to approximately 0.1 mM (Fig. 7T,U). The relation between S100β concentration and [Ca^2+^]_e_ (Fig. 7T) allowed us to estimate the concentration of S100β released by astrocytes causing the 0.13 mM decrease in [Ca^2+^]_e_ recorded following optogenetic activation of astrocytes (see Discussion).

We then examined whether precise control of [Ca^2+^]_e_ induced by applications of S100β or BAPTA of variable duration translated into a precise control of the spiking activity of L5PN (Fig. 7G-N). Indeed, the number of evoked spikes and the firing frequency increased proportionally to the duration of application of S100β (Fig. 7G, I, K, M). The results from 14 neurons were pooled by expressing for each neuron the number of spikes as a percentage of the maximal number of spikes evoked during the strongest spiking response, and the S100β application duration as a function of the maximal duration. The number of evoked spikes (0-100 spikes) increased linearly with the S100β pulse duration (129 µM, 2 to 20 psi, 1-19 s) (R = 0.78, *P* < 0.0001, n = 95 trials in 14 neurons from 11 mice) and a significant correlation was also found between S100β pulse duration and spiking frequency (0-128.5 Hz; R = 0.32, *P* < 0.01). Likewise, gradually increasing the duration of BAPTA pulses (0.5 to 6 s pulses, 10 mM, 1 to 5 psi) also increased the number of spikes and the spiking frequency in L5PN (Fig. 7H, J, L, N). Pooling the data (n = 38 trials from 7 neurons from 6 mice) revealed a linear relationship between BAPTA pulse duration (0.5 to 6 s pulses, 5 or 10 mM, 1 to 10 psi) and the number of spikes (2 to 584 spikes; R = 0.80, *P* < 0.0001), and the spiking frequency in L5PN (0.5-126.3 Hz; R = 0.72, *P* < 0.0001). These data indicate that exogenously applied S100β can finely regulate [Ca^2+^]_e_ and L5PN firing frequency. Decreases of [Ca^2+^]_e_ can affect several conductances that may explain these effects. On the basis of our results with optogenetic stimulation of astrocytes (Fig. 6) and our previous work (Morquette et al. 2015), one likely candidate is I_NaP_.

### Juxtaposition of neuronal Nav 1.6 channels and S100β-positive glial cells in V1

If the effects of S100β are produced by reducing [Ca^2+^]_e_ near voltage-gated sodium channels (Nav) mediating I_NaP_, then astrocytic processes susceptible to release S100β should be closely apposed to these channels. This was examined using immunohistochemistry coupled with confocal and STED microscopy in wild type mice. First, to ensure that S100β-positive processes belonged to astrocytes, we compared localization of S100β to that of GFAP. In all tested mice (n = 5), S100β was present both in the cell body and processes of GFAP-positive cells (Fig. 8D-F). In agreement with previous reports (Caldwell et al. 2000), L5PN expressed Nav 1.6 channels largely on their cell body and apical dendrite (Fig. 8G, I, J-L, N; n = 10 mice). These Nav 1.6-positive L5PN were surrounded by S100β-immunoreactive cells, with S100β-positive processes wrapping around their cell bodies and proximal portions of their dendrites (Fig. 8I-N). This is reminiscent of our observation in GFAP-ChR2-EYFP mice that astrocytic processes positive for S100β and EYFP closely surrounded putative neurons that were S100β-negative and EYFP-negative (asterisk in Fig. 1M,N). Such proximity of S100β and Nav 1.6 channels provides a possible anatomical substrate through which optogenetically-activated astrocytes and their protein S100β evoked Nav 1.6-dependent firing in L5PN.

**Figure 8.**
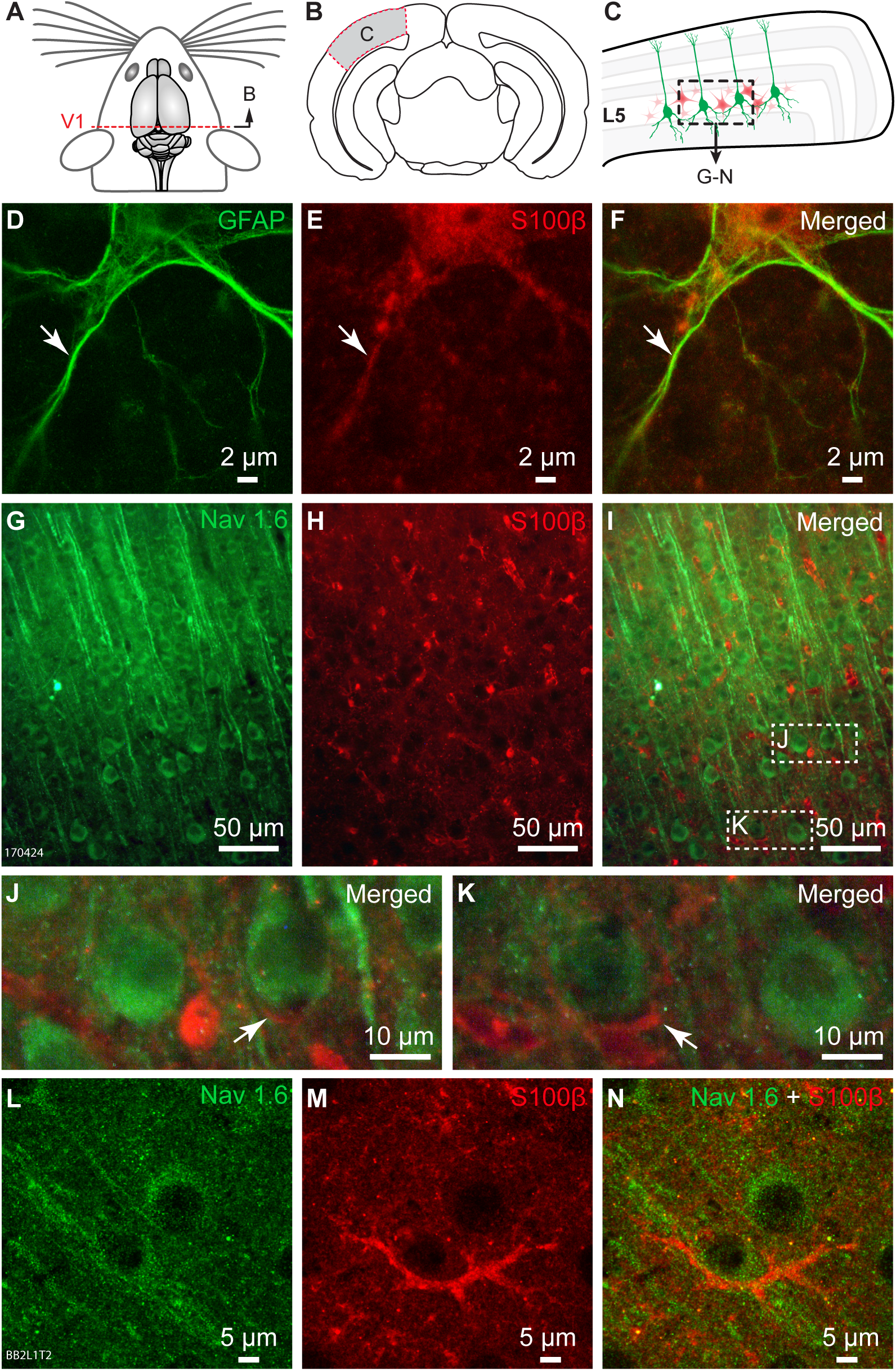
Juxtaposition of Nav 1.6 channels and S100β-positive glial cells. A-B. Dorsal view of the mouse brain showing the level at which the coronal slice of primary visual cortex (V1) shown in B was taken. **C.** Magnification of the grey area in B showing the cortical layers in V1 and, in the black dashed rectangle, the area of layer 5 imaged for Nav1.6-immunoreactive layer 5 pyramidal neurons (L5PN, green) and S100β-immunoreactive astrocytes (red) in panels D-K. **D-F**. STED images showing immunofluorescence against the Glial Fibrillary Acidic Protein (GFAP) (D), S100β (E) and the merged two markers (F). The arrows show a thin astrocytic process showing co-localization of GFAP and S100β. **G.** Immunofluorescence against the Nav 1.6 voltage-gated sodium channels. **H.** Immunofluorescence against the astrocytic Ca^2+^-binding protein S100β. **I.** Merge of the photomicrographs from G-H. **J-K.** Magnifications of the dashed rectangles in I. **L-N.** STED images showing immunofluorescence against the Nav 1.6 voltage-gated sodium channels (L), the astrocytic protein S100β (M) and the merged two markers (N). Data were obtained from 3 different preparations.

### S100β-evoked spiking in L5PN involves Nav1.6 channels

To determine whether S100β exerted its effects directly on L5PN, we first examined whether the spiking activity elicited by S100β local application (2 to 15 s pulses, 2 to 10 psi) was still possible in presence of a cocktail (which did not affect L5PN electrophysiological properties, see Table 2) of bath applied (15-40 min) antagonists targeting AMPA/Kainate (CNQX, 10 µM), NMDA receptors (AP5, 26 µM) and GABA_A_ receptors (gabazine, 20 µM) (14.3 ± 7.1 spikes, n = 4 neurons from 4 mice, 1 to 4 trials per neuron; Fig. 9A-D). Comparison of the activity evoked before and after bath application confirmed that the cocktail did not abolish S100β-evoked spiking in L5PN (control: 26.7 ± 5.4 vs drug: 17.3 ± 9.1 spikes, *P* > 0.05, paired t test, n = 3 neurons from 3 mice).

**Figure 9.**
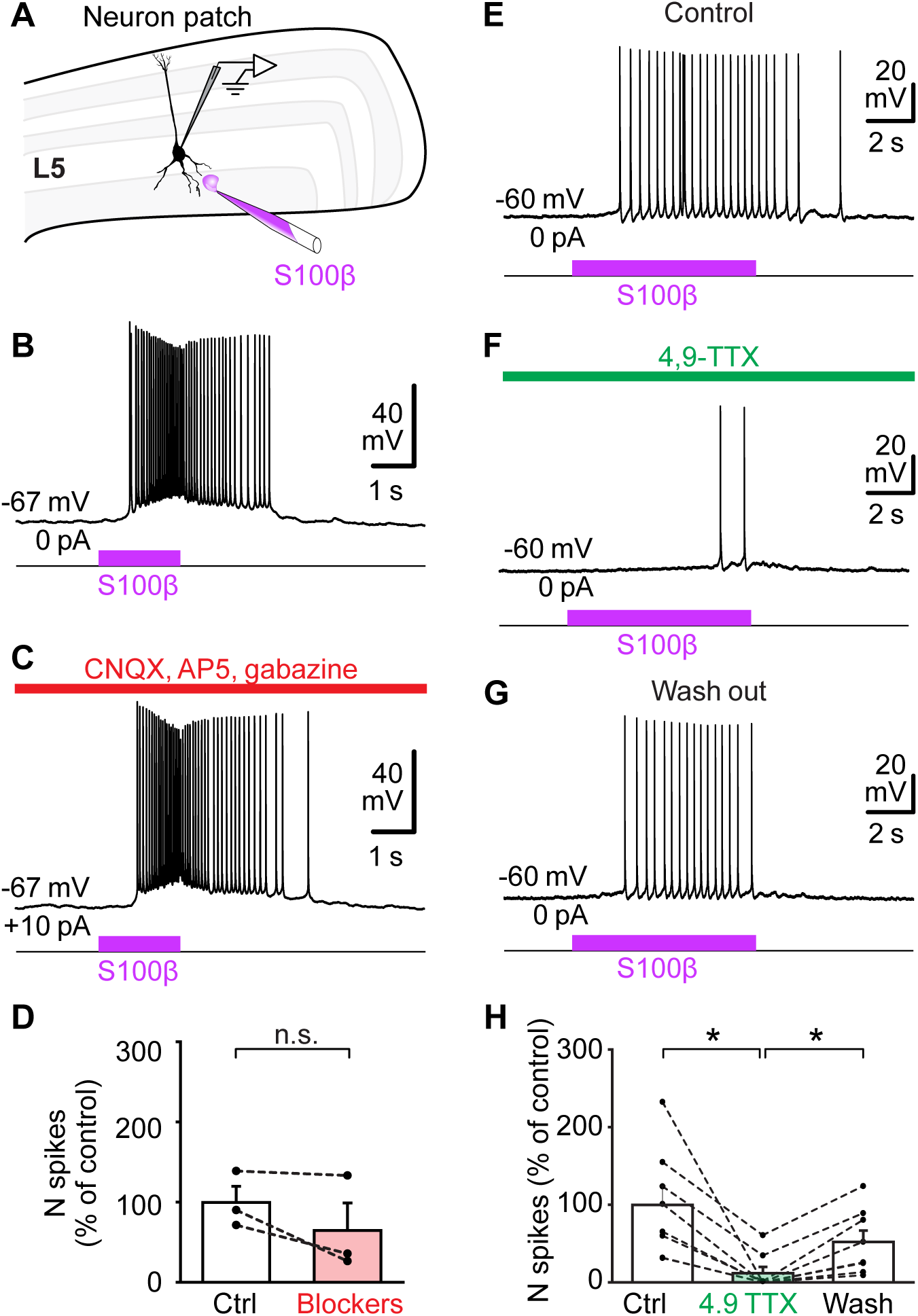
Blockade of Nav 1.6 channels abolishes S100β-evoked spiking in layer 5 pyramidal neurons (L5PN). A. Scheme of the coronal brain slice, where the cortical layers in V1 are indicated. The astrocytic protein S100β was locally applied within layer 5 (L5) over the recorded L5PN. **B-D.** Effect of bath application (40 min) of the fast transmission inhibitors (AMPA/kaïnate receptor antagonist CNQX 10 μM, NMDA receptor antagonist AP5 26 μM, GABA_A_ receptor antagonist gabazine 20 µM) on the spiking activity evoked by local application of S100β (129 µM). In D, effect of the antagonist cocktail on the number (N) of S100β-evoked spikes in 3 neurons, n.s. *P* > 0.05, paired t test. **E-H**. Effect of bath application of the Nav 1.6 channel blocker 4,9-anhydro-TTX (4,9-TTX, 0.1 μM) on spiking activity evoked in L5PN by local application of S100β (129 µM, 5-11 s pulses, 2-10 psi). In H, * *P* < 0.05, Student-Newman Keuls test after a Friedman Repeated Measures Analysis of Variance on Ranks *P* < 0.001. Data from B-C and E-G were obtained from 2 different slices.

We then determined whether S100β-evoked activity involved I_NaP_, as was the case with optogenetic stimulation of astrocytes (Fig. 6). Indeed, bath application (6-26 min) of 4,9-TTX (0.1 µM) (which did not affect L5PN electrophysiological properties, see Table 2) dramatically decreased the spiking activity evoked by S100β (129 µM, 2 to 10 psi, 5 to 11 s pulses) from 30.3 ± 7.4 to 3.7 ± 2.5 spikes (*P* < 0.05 vs. control, Student Newman-Keuls test after a one-way Friedman ANOVA for repeated measures on ranks, *P* < 0.001, n = 8 neurons from 8 slices from 6 mice, 2 to 5 trials per drug condition) (Fig. 9E-F, H). The number of spikes evoked by S100β partially recovered to 15.9 ± 4.5 spikes after washout (*P* < 0.05 vs. drug condition) (Fig. 9G, H).

### S100β overrides the adaptation of firing evoked by L1 stimulation

Next, we examined whether S100β application near L5PN cell body could override the adaptation elicited by trains of stimulations in L1 (Fig. 10) as was shown in Figure 4 with optogenetic activation of astrocytes. Application of 5 s stimulation trains (30-70 µA, 0.2 ms pulses) in L1 at different frequencies (1 to 50 Hz) elicited spiking in L5PN that adapted when frequencies above 10 Hz were used (Fig. 10A-B, E, F). As was the case with optogenetic stimulation of astrocytes (Fig. 4F-H), application of S100β (129 µM, 2 to 10 psi, 5 s pulses) near the recorded L5PN cell body prevented firing adaptation at higher stimulation frequencies delivered to L1 (Fig. 10C) as shown by measures of spiking response duration (Fig. 10E, for 10, 20, 30, 40, and 50 Hz, *P* < 0.05 to *P* < 0.001 vs. control, Student-Newman-Keuls test after a two-way ANOVA for repeated measures on ranks, *P* < 0.001) and of the number of spikes (Fig. 10F, for 1, 20, 30, 40, and 50 Hz, *P* < 0.05 to *P* < 0.001 vs. control, Student-Newman-Keuls test after a two-way ANOVA for repeated measures on ranks, *P* < 0.01, n = 5 neurons from 5 slices from 5 mice). These effects were transitory and disappeared after S100β was washed out (Fig. 10D) both for spiking response duration (Fig. 10E, for 30, 40, 50 Hz, *P* < 0.001 vs. application of S100β) and for the number of spikes (Fig. 10F; for 1, 30, 40, and 50 Hz, *P* < 0.05 to *P* < 0.001 vs. application of S100β). These results show that application of exogenous S100β was sufficient to profoundly modify the ability of L5PN to process inputs from distal dendrites, as was the case with optogenetic activation of astrocytes (Fig. 4).

**Figure 10.**
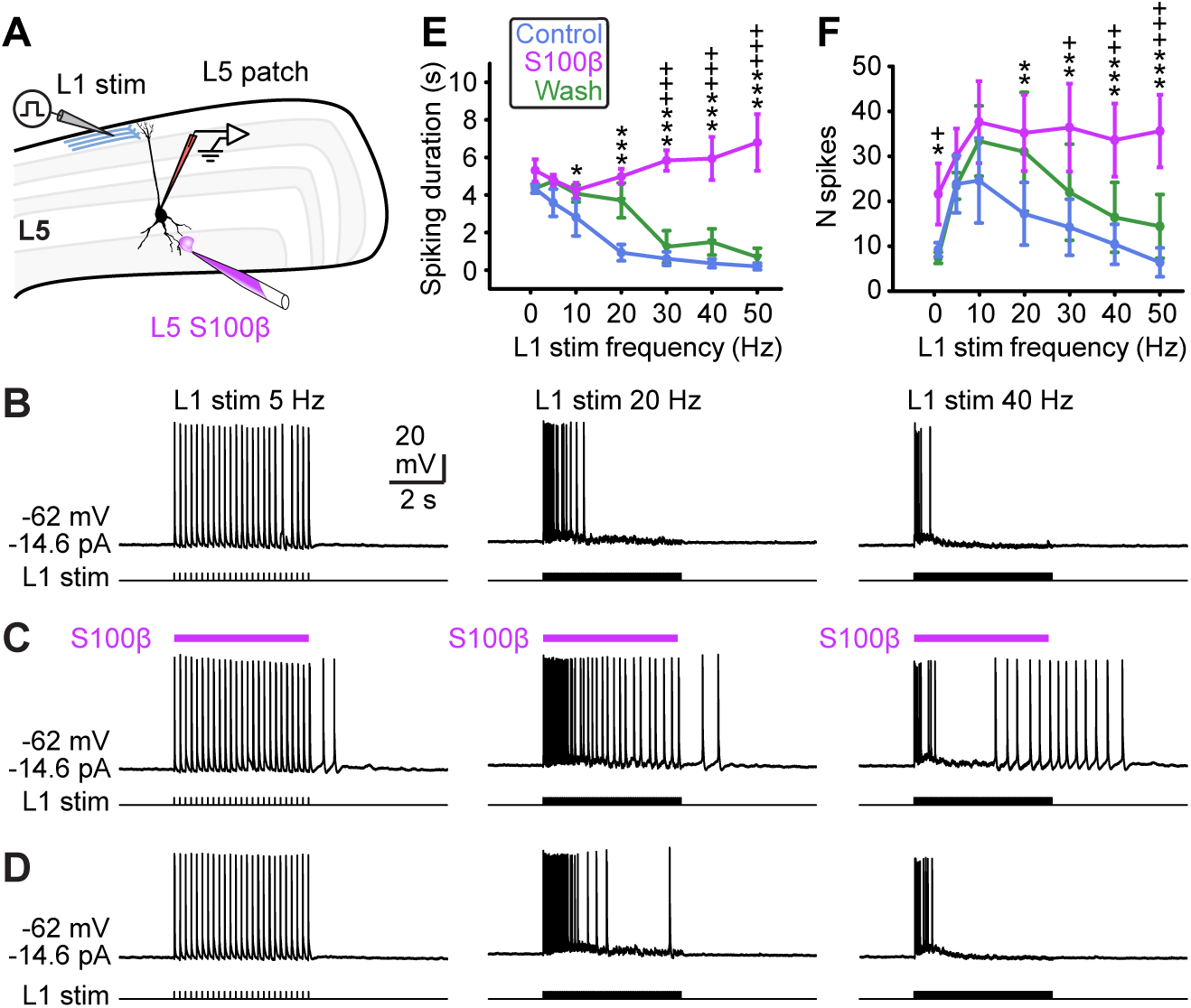
The astrocytic protein S100β applied in layer 5 (L5) overrides the adaptation of layer 5 pyramidal neuron (L5PN) firing evoked by repetitive L1 stimulation. A. Scheme of the V1 coronal brain slice, showing positioning of the pipette for L5 local drug application and of the recording and stimulating electrodes in L5 and L1 respectively. **B-D.** The responses evoked by electrical stimulation of L1 axons (at 1, 5, 10, 20, 30, 40, 50 Hz, 0.2 ms pulses, 5 s train, 30-70 µA) contacting the recorded L5PN were investigated in control condition (B), during local application of S100β (129 µM, 5 s pulses, 5 psi) near the recorded L5PN (C), and after wash-out (D). **E.** The duration of the spiking response was compared for each L1 stimulation frequency (\**P* < 0.05, \*\*\**P* < 0.001 vs. control; ^+++^*P* < 0.001 vs. drug condition, Student-Newman-Keuls test after a two-way ANOVA for repeated measures on ranks *P* < 0.001, pooled data from 5 neurons from 5 slices from 5 mice). **F.** The number of spikes were compared for each L1 stimulation frequency (\**P* < 0.05, \*\**P* < 0.01 vs. control, \*\*\**P* < 0.001 vs. control; ^+^*P* < 0.05, ^++^*P* < 0.01, ^+++^*P* < 0.001 vs. S100β condition, Student-Newman-Keuls test after a two-way ANOVA for repeated measures on ranks *P* < 0.01, pooled data from 5 neurons from 5 slices from 5 mice).

## Discussion

The results presented here provide evidence that cortical astrocytes can induce spiking in L5PN and alter their input-output transfer function by potentiating their I_NaP_, through an S100β-mediated decrease of [Ca^2+^]_e_ (Fig. 11).

**Figure 11.**
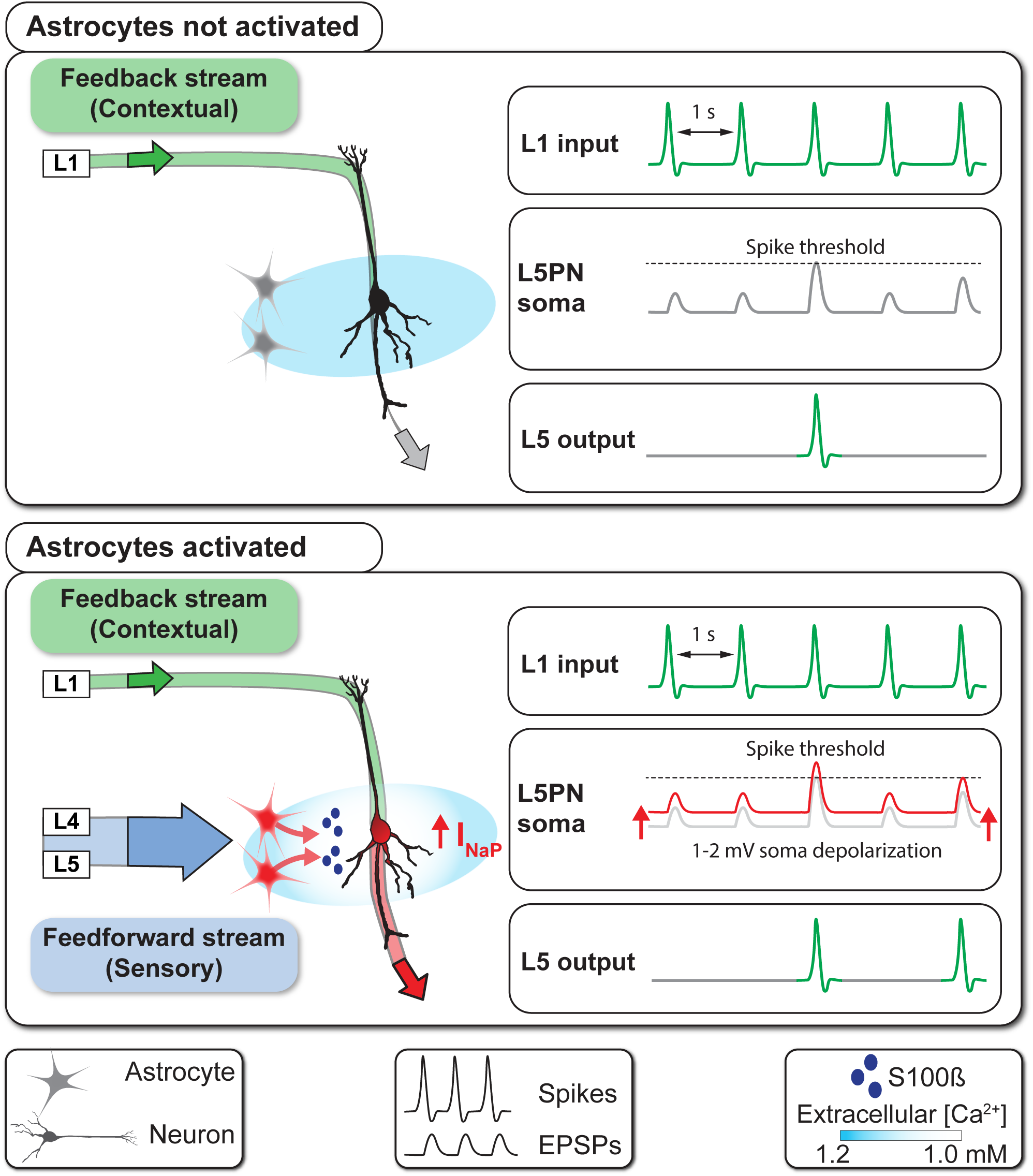
Model of the S100β-dependent mechanism through which astrocyte control neural firing in cortical circuits. Layer 5 pyramidal neurons (L5PN) receive a stream of feedback inputs on the distal tuft of their apical dendrite in layer 1 (L1) (green arrow in L1) and feedforward stream of inputs on their basal dendrites proximal to the soma (blue arrow in L4-L5 of lower panel). Distal inputs (green traces in L1 input box, upper and lower panels) do not always bring the neuron to firing threshold (grey traces under the dashed line in L5PN soma box). Only those reaching threshold will trigger spiking (green trace in L5 output box). Activation of astrocytes (lower panel) by proximal inputs leads to release of the calcium (Ca^2+^)-binding protein S100β (blue dots) and subsequent decrease of extracellular Ca^2+^ which in turn potentiates Nav 1.6 channel activation and further depolarizes the neuron (shift from grey to red traces in L5PN soma box of lower panel) enabling subthreshold inputs to reach threshold and initiate a spike. Through this mechanism, astrocytes can control spiking frequency and the integration of incoming inputs from L1, and thus contribute to two core units of neural computations.

### Technical considerations

Several studies have reported that GFAP can be expressed in some neurons during development (Casper and McCarthy 2006; Fujita et al. 2014; Ganat et al. 2006; Platel et al. 2009; Su et al. 2004). Thus, the use of a GFAP promoter to drive expression of ChR2 can raise concerns about specificity of expression in astrocytes. However, although they may not suffice to completely rule out the expression of ChR2-EYFP in neurons in our study, the following observations argue against the possibility that this may be the main factor for the effects reported here: 1) The vast majority (94%) of V1 neuronal cell bodies or processes labeled with NeuN or MAP2 immunofluorescence, or with biocytin following patch-clamp experiment, were found to be negative for EYFP, while on the contrary S100β-positive astrocytes expressed EYFP in our ChR2-EYFP mice; 2) Effects of photo-stimulation were observed in a very large sample of neurons in the present study, whereas levels of expression of GFAP in neurons, although variable among studies, are quite low in most of them (as low as 0.3% of cortical neurons according to Ganat et al. 2006), making it very unlikely that we recorded in most cases the rare cortical neurons that express or have expressed GFAP during their lifetime; 3) Responses elicited by photostimulation persisted in presence of glutamatergic, GABAergic and purinergic receptors blockers, ruling out the possibility of transmitter release from neuronal fibers expressing ChR2-EYFP. In line with this,MAP2-positive fibers were EYFP-negative; 4) Spiking responses elicited by optogenetic stimulation were nonetheless abolished by the anti-S100β antibody; 5) Although rapid, the optogenetically evoked neuronal response in our largest sample (55 neurons for 20 s long photo-stimulations) occurred on average 1.42 s after beginning of the light pulse, which is much slower than responses elicited by expression of ChR2 in neurons.

The second technical limitation in our study is the use of ChR2 to stimulate astrocytes. ChR2 has been associated with changes in pH in astrocytes (Losi et al. 2017) and a recent study by Octeau (2019), showed that its optogenetic excitation leads to transient increases in extracellular K^+^ raising the possibility that the firing that persisted in presence of the antagonists could have also resulted from accumulation of extracellular K^+^. However, such nonspecific effects are unlikely to be targeted by the anti-S100β antibody, which should not in principle affect pH or the K^+^ increase.

It remains to be confirmed whether the effects reported here can be reproduced in adults since our results were obtained in P11-P31 mice, right before the chemical synapses stabilize and the astrocytes acquire their mature morphology in the cortex (Allen and Eroglu 2017; Farhy-Tselnicker and Allen 2018). However, a recent study showed in adult rodents that cognitive flexibility, an important function of the medial prefrontal cortex which relies on neuronal oscillations and coupling across frequency ranges is improved by injections of S100β or chemogenetic activation of astrocytes and impaired by inactivation of endogenous S100β (with an antibody as we did here and previously) or by a reduction in astrocyte number (Brockett et al. 2018), suggesting that S100β-dependent control of neural activity is maintained during adulthood.

### Impact of [Ca^2+^]_e_ fluctuations on cell signalling

Decreases of [Ca^2+^]_e_, varying from 0.1 to 0.6 mM and lasting up to tens of seconds, have been reported in cortical and cerebellar circuits following low frequency (20 Hz) stimulation, periods of intense activity, or with changes in the sleep/wake cycle (Amzica et al. 2002; Ding et al. 2016; Nicholson et al. 1978). There is ample evidence suggesting that such variations in [Ca^2+^]_e_ can serve for cell signalling in the brain (for review, see MacDonald et al. 2006). The decreases in [Ca^2+^]_e_ observed here following optogenetic stimulation of astrocytes (of 0.13 mM in average) are of the same order of magnitude as some of the above studies (Ding et al. 2016; Nicholson et al. 1978). However, they probably represent an underestimation of the effective local reduction in [Ca^2+^]_e_ because it is difficult to position the tip of Ca^2+^-sensitive electrodes near the precise location where the Ca^2+^ decreases are occurring, and because the recordings were obtained from submerged preparations that were constantly perfused with a solution containing a constant [Ca^2+^]_e_. A decrease in [Ca^2+^]_e_ can affect neural activity in multiple ways. Many ionic channels are affected by membrane electric fields established by interactions of divalent cations with fixed anions. Thus, changes in divalent ion concentrations alter these fields and lead to what is called screen charge effects (Hille 2001; Jones and Smith 2016), but beyond these effects, changes in [Ca^2+^]_e_ of sometimes as little as 0.1 mM, have been reported to directly affect several channels and transporters (for review, see Jones and Smith 2016) including voltage-gated sodium channels (Nav) (Armstrong and Cota 1991). Our observation that the effects induced by S100β local application or optogenetic stimulation of astrocytes were nearly abolished by 4,9-TTX, a selective blocker of Nav1.6 (Hargus et al. 2013; Rosker et al. 2007) strongly suggests that spiking resulted from an action of Ca^2+^ on Nav1.6 channels. This is in line with previous reports showing that Ca^2+^ changes the kinetics or stability of Nav channels by interacting either directly with extracellular moieties (Armstrong and Cota 1990), amino acid residues lining the channel’s pore (Santarelli et al. 2007), or indirectly with calmodulin or calpain that respectively modify the gating behaviors of the channel (Sarhan et al. 2012) or cleave Nav channel α subunit Nav1.6 (Brocard et al. 2016).

We, and others, have already shown that decreases of [Ca^2+^]_e_ are capable to shift the activation threshold of I_NaP_ and its half-activation voltage (by ∼1 mV, every 0.1 mM; Brocard et al. 2013) towards more hyperpolarized potentials (Li and Hatton 1996; Morquette et al. 2015; Su et al. 2001). Thus, even small variations in [Ca^2+^]_e_ can have an important effect on the neuron’s firing by decreasing the threshold for I_NaP_ activation and increasing its amplitude. Our previous work suggested that S100β can significantly reduce [Ca^2+^]_e_ (Morquette et al. 2015), and raised the important question of whether sufficient amounts of S100β can be released from astrocytes to achieve this effect on I_NaP_.

### Can endogenous S100β induce sufficient fluctuations of Ca^2+^ in situ?

S100β is detected in the extracellular space in multiple species and conditions (Shashoua et al. 1984; Van Eldik and Zimmer 1987; Vicente et al. 2007, for review see Donato et al. 2009) and its level in the serum or cerebrospinal fluid are known to increase under various neuropathological conditions including Alzheimer’s disease, epilepsy, or schizophrenia (Griffin et al. 1989, 1995; Rothermundt et al. 2004; Schmitt et al. 2005). Many factors, including neuronal activity and increases in astrocytic cytosolic Ca^2+^, are known to increase S100β release, but the mechanism underlying this release remains ill-defined, although mGluR3 and/or A1 adenosine receptors, but not connexin 43 hemichannels, may be involved under some circumstances (Ciccarelli et al. 1999; Sakatani et al. 2008). It is unknown whether the protein is *secreted* through pores (or channels) or *released* from vesicles that can concentrate it before release. In Schwann cells and glioblastoma, endogenous S100β is localized in vesicle-like structures (Davey et al. 2001; Perrone et al. 2008) that are translocated across the cell in response to increasing intracellular Ca^2+^ or decreasing intracellular Zn^2+^ concentration through an endoplasmic reticulum-Golgi-independent pathway (Davey et al. 2001) suggesting that it could be released from vesicles. Additional support is provided by the observation that extracellular S100β is recaptured in a time-dependent manner by vesicle endocytosis (Lasič et al. 2016). The exact concentration of the protein at the site of release is difficult to estimate, but on the basis of our measurements of S100β-evoked decrease in [Ca^2+^]_e_ (Fig. 7T), we estimated that the 0.13 mM decrease in [Ca^2+^]_e_ evoked by optogenetic stimulation of astrocytes (Fig. 6C) would correspond to a pulse of 233 ms at 10 psi, which was estimated to contain 0.62 ± 0.35 fmol of S100β, i.e. 6.7 ± 3.8 pg of S100β (see Methods). Such quantity is compatible with previous measurements in hippocampal brain slices where the extracellular concentration of S100β was found to increase 5 fold (from 49 to 240 ng/ml) after a 30 min exposure to 400 nM of kainate (Sakatani et al. 2008), which corresponds to an increase from 11 to 54 pg of S100β for an estimated extracellular volume of 0.225 µL (i.e. 15% of a 1.5 µL hippocampal brain slice, see Sakatani et al. 2008). Thus, considering how the small extracellular volume of the brain restricts diffusion (slowing it by a factor of five according to Kullmann et al. 1999) and given the close proximity between S100β-positive astrocytic processes and Nav1.6 channels observed here using immunohistochemistry (Fig. 8N), it is reasonable to assume that the activation of astrocytes can translate into the release of sufficient amounts of S100β to produce a detectable effect on [Ca^2+^]_e_.

### Conditions to elicit astrocytic control of L5PN output

To play a key role in brain physiology, the proposed mechanism, which we assume to be triggered by increases in astrocytic Ca^2+^ signalling, needs to be spatially and temporally fine-tuned for the proper control of L5PN output that is largely determined by either slow, widespread inputs from neuromodulators related to vigilance states, or fast, spatially constrained inputs, related to sensory information processing (in the visual cortex). The levels of noradrenaline and acetylcholine vary with vigilance and attention states, and these neuromodulators can induce large transients in astrocytic cytosolic and microdomains Ca^2+^ signals (Agarwal et al. 2017; Chen et al. 2012; Ding et al. 2013; Paukert et al. 2014; Srinivasan et al. 2015; Takata et al. 2011). These slow signals, which resemble those we imaged in astrocyte cell bodies following optogenetic stimulation (Fig. 2E) could potentially lead to release of S100β and explain the extracellular Ca^2+^ fluctuations observed during the sleep/wake cycle (Ding et al. 2016). Our data provide evidence that the S100β-mediated astrocytic mechanism produces sustained effects on input integration or spiking activity during seconds to dozens of seconds, a time scale compatible with a classical slow neuromodulatory effect on behavior.

Astrocytic Ca^2+^ signals triggered by sensory stimulation have also been reported in many studies (Asada et al. 2015; Gee et al. 2014; Sonoda et al. 2018; Takata et al. 2011; Wang et al. 2006). It has been debated whether these astrocytic responses result from glutamate released from thalamocortical fibers and intracortical circuitry or from noradrenaline released from ascending fibers of Locus Coeruleus neurons that are activated by sensory inputs (Ding et al. 2013; Paukert et al. 2014; Srinivasan et al. 2015). However, the sensory-evoked astrocytic activation pattern is often very constrained spatially and reflects somatotopy and/or physical attributes of the stimulus used (for review see López-Hidalgo and Schummers 2014), like is the case in the ferret visual cortex where astrocytic responses have sharper tuning curves than surrounding neurons (Schummers et al. 2008). Such sensory-evoked responses show a certain degree of specificity relative to the stimulated part of the body, arguing against a “general” widespread activation through noradrenergic inputs. For instance, hind paw stimulation has no effect on astrocytic Ca^2+^ signaling in brain astrocytes responding to whisker stimulation and vice versa (Stobart et al. 2018b).

### Timescales of astrocytic and neuronal activities

Recent observations uncovered that astrocyte signals occur in parallel at different time scales following sensory stimulation. Drugs that interfere with noradrenergic and cholinergic transmission reduce the large astrocytic Ca^2+^ responses that follow sensory stimulation, but unmask smaller sensory related responses (Sonoda et al. 2018). Interestingly, the large astrocytic responses reduced by these drugs are those occurring with a delay while the timing and amplitude of fast responses observed in microdomains are not altered (Stobart et al. 2018a). These fast events display a wide range of latencies when compared to neuronal Ca^2+^ signals that occur simultaneously, but a fraction (8%) of astrocytic signals (detected only in microdomains with the membrane-tethered genetically encoded Ca^2+^ indicator Lck-GCaMP6f) had an onset latency (median onset = 0.588 s) that did not differ from that of neurons (Stobart et al. 2018a). This, combined to the short latencies that we observed for neuronal response (median = 0.853 s, n = 55 trials from 55 neurons) following the onset of optogenetic stimulation of astrocytes, suggest that sensory inputs can evoke fast and localized astrocytic Ca^2+^ increases, which in turn could trigger a rapid, specific and spatially constrained S100β-mediated modulation of neuronal function.

### Involvement of other gliotransmitters

Astrocytes are known to influence neural activity through regulation of extracellular K^+^ ions and release of glutamate (Bezzi et al. 1998; Parpura et al. 1994; Poskanzer and Yuste 2011, 2016), GABA (Kozlov et al. 2006; Liu et al. 2000), ATP (Newman 2003; Panatier et al. 2006; Poskanzer and Yuste 2011, 2016) or the NMDA co-agonist D-Serine (Henneberger et al. 2010; Yang et al. 2003). On the basis of these and other studies (e.g. Torres et al. 2012), we expected to have large effects when blocking glutamatergic, GABAergic and purinergic receptors. Indeed, antagonists of fast ionotropic glutamatergic and GABAergic receptors reduced the effects of optogenetic activation of astrocytes. Unexpectedly, these effects were counteracted by addition of metabotropic glutamatergic receptor blockers. This may be explained by the fact that metabotropic receptors were reported to be located mostly on terminals of inhibitory interneurons. Suramin produced mixed effects, and no clear statistical difference was found. These mixed effects may have masked or counteracted each other when considering the effects of all blockers, but nonetheless, the anti-S100β antibody induced the strongest blockade of the neural responses evoked by astrocyte optogenetic activation. The unexpected lack of strong blockade of the neural response with all receptors antagonists may result from the fact that we stimulated only small precise areas in L5 close to the recorded L5PN, and such areas may not involve the exact same circuitry as those examined in other studies conducted in layers 2/3 or in the hippocampus. Another reason may be that the effect reported here rests on an ionic conductance (I_NaP_) that is modulated by [Ca^2+^]_e,_ which is not necessarily the case for other studies. Torres et al. (2012) considered the effects of [Ca^2+^]_e_ decreases and documented ATP mediated effects on inhibitory interneurons. However, their study was conducted in the hippocampus, and importantly, used a [Ca^2+^]_e_ of 2mM in the aCSF, which may explain the difference with our results. Here we used a physiological [Ca^2+^]_e_ (Jones and Keep 1988) and observed effects on firing only when [Ca^2+^]_e_ dropped to ∼1.1 mM, a condition never reached in their study.

### Significance for L5PN function

L5PN are associative neurons, which integrate sensory inputs (feedforward stream in Fig. 11) to their proximal dendrites with information from other parts of the cortex arriving at their distal apical dendritic tuft (Larkum 2013). These associative properties are believed to confer to these neurons the ability to associate information from the sensory inputs with an internal representation of the world (feedback stream in Fig. 11) in order to make predictions (Felleman and Van Essen 1991; Larkum 2013). We show that activation of proximal astrocytes within L5 potentiates the integration of inputs to distal apical dendritic tuft by depolarizing the soma of L5PN. Moreover, our observation that astrocytic activation overrides L5PN firing adaptation induced at higher stimulation frequencies of L1, combined to the fact that astrocytic response to sensory stimulation occurs over a longer period than neurons (Stobart et al. 2018a), expands the time window during which distal and proximal inputs can interact, and by doing so astrocytes are in good position to have a direct impact on cognitive functions that depend on the associative ability of L5PN.

The APs generated in L5PN soma by astrocyte activation could backpropagate in the apical dendrites and potentiate the ability of the distal inputs to influence L5PN firing. When backpropagating AP coincide with distal dendritic inputs, a dendritic Ca^2+^ spikes is generated, and this translates in a burst of AP in the soma characterized by high intraburst firing frequencies (Larkum et al. 1999) that resemble those evoked here by S100β (Fig. 7E), BAPTA (Fig. 7F) and optogenetic activation of astrocytes (Fig. 3F).

### Conclusions

Our study brings a new dimension to the mechanisms by which astrocytes control firing and input integration, two essential components of brain computations. Our work suggests that the localized action of S100β in controlling [Ca^2+^]_e_ and consequently Nav1.6-mediated induction of L5PN spiking could be a powerful control mechanism of cortical associations. The control exerted by astrocytes on neuronal activity through the regulation of the extracellular ionic environment is increasingly considered (Kadala et al. 2015, see also Ashhad and Narayanan 2018). Here we have focused on how astrocytes modulate L5PN output, but effects on other types of neurons should be considered as well since stimulation of astrocytes has been shown to modulate orientation selectivity of excitatory but also inhibitory interneurons in the visual cortex (Perea et al. 2014), adding a potential new layer of complexity to this signalling mechanism. Further, our work demonstrates that such S100β-dependent mechanism, which was first found to control neural activity in brainstem motor circuits (Morquette et al. 2015), is likely to be widespread in the brain. The mechanism we report here is likely involved in the effects on cognitive flexibility, reported in adult mammals following intracortical injections of S100β, chemogenetic activation of astrocytes, and inactivation of endogenous S100β (Brockett et al. 2018). Future work should determine in which conditions astrocytes are made to release S100β in the extracellular space and how this mechanism may be regulated because its dysfunction could translate into abnormal levels of S100β in the extracellular space, leading to abnormal levels of [Ca^2+^]_e_, and undesirable effects on neuronal activity, which may explain why several brain disorders are associated to elevated S100β serum and cerebrospinal fluid levels.

## Acknowledgments

We thank S. Miranda-Rottmann for help genotyping the mice, R. Robitaille for the gift of the SC-71 antibody, J. Lainé for technical help with the Leica nanoscope, and D. Veilleux for technical assistance. This work was supported by the Canadian Institutes of Health Research (14932 to A.K., 407083 to D.R.); Natural Sciences and Engineering Research Council of Canada (RGPIN-2017-05522 and RTI-2019-00628 to D.R.), the Fonds de la Recherche en Santé du Québec Junior 1 fellowship (34920 and 36772 to D.R.).

## Author contributions

D.R., M.H.D., R.A., and A.K. designed research. D.R., M.H.D., S.C, B.J.B, M.F. performed research. D.R., M.H.D., B.J.B, S.C, M.F. and A.K. analyzed data. D.R., R.A., and A.K. wrote the paper.

## Competing interests

The authors have no competing interest to declare.

## Notes

### Competing Interest Statement

The authors have declared no competing interest.

